# Heterogeneity and evolution of DNA mutation rates in microsatellite stable colorectal cancer

**DOI:** 10.1101/2024.02.26.582054

**Authors:** Elena Grassi, Valentina Vurchio, George D. Cresswell, Irene Catalano, Barbara Lupo, Francesco Sassi, Francesco Galimi, Sofia Borgato, Martina Ferri, Marco Viviani, Simone Pompei, Gianvito Urgese, Bingjie Chen, Eugenia R. Zanella, Francesca Cottino, Alberto Bardelli, Marco Cosentino Lagomarsino, Andrea Sottoriva, Livio Trusolino, Andrea Bertotti

## Abstract

DNA sequence mutability in tumors with chromosomal instability is conventionally believed to remain uniform, constant, and low, based on the assumption that further mutational accrual in a context of marked aneuploidy is evolutionarily disadvantageous. However, this concept lacks robust experimental verification. We adapted the principles of mutation accumulation experiments, traditionally performed in lower organisms, to clonal populations of patient-derived tumoroids and empirically measured the spontaneous rates of accumulation of new DNA sequence variations in seven chromosomally unstable, microsatellite stable colorectal cancers (CRCs) and one microsatellite unstable CRC. Our findings revealed heterogeneous mutation rates (MRs) across different tumors, with variations in magnitude within microsatellite stable tumors as prominent as those distinguishing them from microsatellite unstable tumors. Moreover, comparative assessment of microsatellite stable primary tumors and matched synchronous metastases consistently documented a pattern of MR intensification during tumor progression. Therefore, wide-range diversity and progression-associated evolvability of DNA sequence mutational instability emerge as unforeseen hallmarks of microsatellite stable CRC, complementing karyotype alterations as selectable traits to increase genetic variation.

**One sentence summary:** Tumors with chromosomal instability accrue DNA sequence mutations at highly variable rates, which increase during metastatic progression.

## Introduction

Genetic instability, whether in the form of DNA hypermutability or karyotypic aberrations, bestows cancer cells with the ability to accumulate new genetic variants at an accelerated pace (*1-6*). In turn, an increased mutation rate encourages the diversification of evolving clades of cells that compete following Darwinian rules, thereby ensuring sufficient genetic heterogeneity to overcome selection barriers such as apoptosis, senescence and therapeutic pressure (*2*).

Although genetic instability is universally acknowledged as fundamental to cancer onset and progression and a key contributor to therapy resistance (*7*), an empirical and quantitative approach for reliable MR estimation in human tumors is still lacking (*8-11*). Several computational efforts have been made to infer tumors’ MRs from bulk DNA sequencing data (*12,13*), based on the assumption that the cumulative distribution of subclonal variant allele frequencies (VAFs) changes as a function of MR (*14*). While these *in silico* analyses have indicated potential MR heterogeneity across tumors (*10*), none of these predictions have been experimentally validated or benchmarked against a known ground truth. Some wet lab approaches have also been pursued by customizing MA experiments typically conducted in lower unicellular species, where lines of organisms are propagated by single-progeny descent to compute the differential repertoire of mutations arising over a given number of cell divisions (*15*). However, these attempts have relied on immortalized cancer cell lines, long adapted to grow on plastic and likely undergoing evolutionary dynamics that may have altered the original tumor’s MR (*16*); more importantly, MA assays in cell lines were limited to small datasets, inadequately sized to measure MR variance across individuals (*9*).

Colorectal cancer (CRC) serves as a prototypical example of neoplastic disease propelled by genetic instability. On one side, a subset of CRC tumors exhibit microsatellite instability (MSI), marked by the accumulation of single-base and indel mutations altering the length of tandem repeats within microsatellite regions due to defects in the DNA mismatch repair system (*17*). On the other side, the majority of CRCs display chromosomal instability (CIN), leading to gross chromosomal rearrangements and aneuploidy (*17*); it is commonly presumed that CRC CIN tumors have uniformly low and constant MRs at the DNA sequence level, with genetic diversity primarily fueled by structural genomic abnormalities. To challenge these assumptions within an experimentally testable setting, applied to clinically relevant model systems, we tailored classical MA methods and estimated MR values in a representative cohort of patient-derived CRC tumoroids.

### DNA mutation rate heterogeneity in MSS CRC

To gauge MR variability in CRC, we took advantage of a collection of over 100 patient-erived tumoroids (PDTs) available at our institute (*18*). From this pool, we selected a subset of eight distinct PDTs that recapitulated some of the most prevalent molecular variants in CRC, including one KRAS mutant MSI tumor and seven CIN, microsatellite stable (MSS) lesions (among the latter, three harbored KRAS mutations and one exhibited a TP53 wild-type status) (Table S1). The seven MSS PDTs were obtained from five patients: two originated from a synchronous metastasis and the corresponding primary tumor; another pair originated from metachronous metastatic lesions; the remaining three MSS samples originated from independent metastatic lesions from as many patients.

To reliably measure the MR of each PDT, we performed a MA assay (*15*) employing a sequential single-cell cloning approach (Fig. 1A). In MA experiments, a clonally expanding cellular population is subjected to repeated bottlenecks, thus minimizing the impact of natural selection and facilitating divergence through the accumulation of neutral mutations by random genetic drift. By comparing the accumulated mutations between the endpoint and ancestor of a lineage and adjusting for the number of cell generations, the mutation rates of various types of DNA alterations, ranging from point mutations to aneuploidies, can be estimated. In our experimental setting, each PDT was single-cell cloned in multiple replicates (n = 2-4, referred to hereafter as T0 clones), and cloned lineages were passaged repeatedly to allow for the accumulation of *de novo* mutations. To maintain conditions conducive to neutral evolution (*19*), we periodically induced bottlenecks by dissociating clones and replating approximately 100 random individual cells every two weeks (see Methods for details). After six months, single-cell subclones (n = 1-6, referred to as T1 subclones) were derived from each original clone. All T0 clones and T1 subclones (n = 97) underwent whole-genome sequencing (WGS) shortly (∼6 weeks) after derivation to obtain a high-quality proxy of the genome of the founder cell at the time of isolation (*20*) (Table S2).

**Figure 1.**
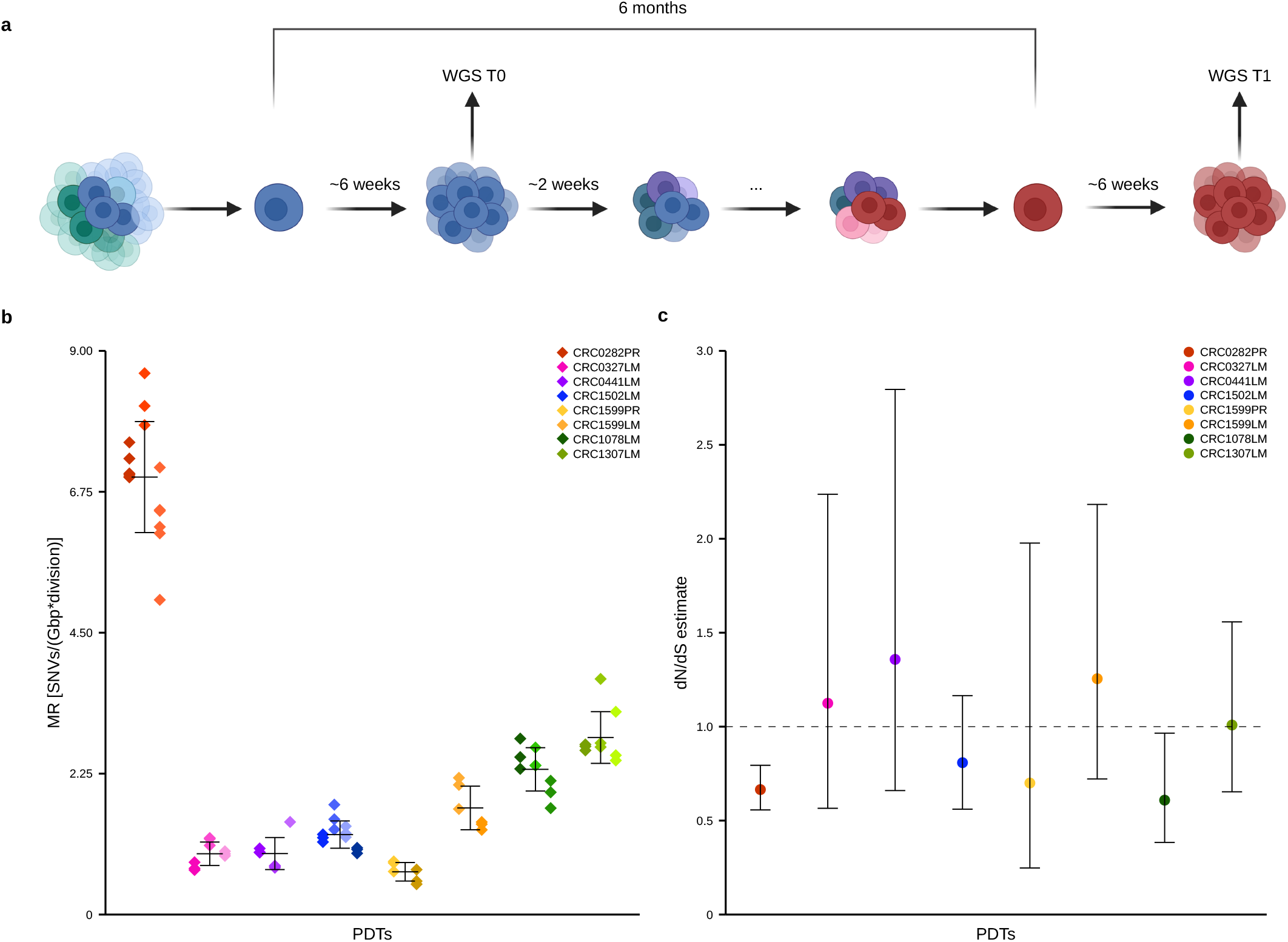
DNA mutation rates across different CRC PDTs. **a**, Schematic overview of the MA experimental design illustrating the repeated passaging of single-cell derived clones over a six-month period, with 100-cell bottlenecks every two weeks. Created with BioRender.com. **b**, SNVs mutation rates in PDTs. Each dot represents the MR estimate of an individual T1 subclone (*N* = 6-15 for each PDT; total *N* = 73); colors correspond to the parental PDT (*N* = 8), while shades of colors indicate the T0 clone from which each T1 subclone originated. *P* = 7.13e-12, Kruskal-Wallis rank sum test. CRC0282PR vs CRC1307LM, *P* = 1.53e-06; CRC1307LM vs CRC1599PR, *P* = 0.0004, Wilcoxon rank sum test. Error bars indicate the average ±1 standard deviation. **c**, dN/dS estimates for the whole set of *de novo* variants accumulated during the MA experiment. The maximum likelihood estimate of each PDT, obtained using the statistical model dNdScv, is represented as a colored dot. Error bars indicate the upper and lower bounds of the inferred dN/dS values.

We normalized the number of mutations gained at T1 against those found in T0 clones for the length of the analyzed genome. To avoid sensitivity biases, the analysis was restricted to genomic regions with a copy number ranging between 1 and 3 in both T0 clones and T1 subclones (see Methods and Table S3). The number of DNA replications was calculated using EdU staining as an estimate of new DNA synthesis (see Methods and Fig. S1). The ratio of the number of newly acquired mutations to the number of DNA replications during the six-month time period between T0 and T1 yielded an MR value for each clone (Fig 1B and Table S3).

We observed a similar MR in all the clones derived from the same tumor, with every PDT showing a distinct mutability footprint discernibly separate from that of the other models. As expected, the MSI tumor (CRC0282PR) was characterized by a particularly elevated MR, with a 2.4-fold increase compared to the MSS tumor with the highest MR (CRC1307LM) when considering the accumulation of single-nucleotide variants (SNVs) (Fig. 1B) and a 17-fold increase when evaluating InDel accumulation (Fig. S2 and Table S3). We note that MSS tumors had a magnitude of MR variation comparable to (and even higher than) that between the MSI and the MR-high MSS PDTs, with a 4.1-fold MR difference between the highest and lowest MSS models in terms of SNV accrual (Fig. 1B and Table S3). This indicates that the MRs of MSS tumors are heterogeneous, and the dynamics of SNV *de novo* generation can vary across MSS tumors to a similar extent as they differ between MSI and MSS samples.

These findings were robust to methodological biases as they were consistent when computing the MR using cell-population doublings instead of EdU incorporation to estimate the number of cell divisions (Fig. S3A). Moreover, both inter-model variability and intra-model consistency were observed when examining the absolute number of gained mutations (Fig. S3B). We also note that, although subject to some degree of uncertainty, the MR estimates based on our experiments are quantitative and not relative, providing measures that could be used to benchmark MR estimates based on the interrogation of cancer genomic datasets.

The dN/dS analysis confirmed that the accumulation of new mutations occurred in a context of neutral growth, with a subtle trend towards negative selection that reached significance in two of the tumoroid models (Fig 1C). This slight deviation can be reasonably explained by the repetitive 100-cell bottlenecks implemented to maximize neutrality, which likely did not entirely prevent the counterselection of heavily deleterious mutations severely hindering cell proliferation or triggering cell death (*19*). The distribution of *de novo* mutations across the genome mirrored that of truncal variants in parental (pre-cloning) PDTs, with a marginal decrease in variants acquired within coding regions (3.8% versus 3.9%) and a modest increase in variants located within introns (48.4% versus 47.9%). This finding is consistent with the absence of positive selection during the MA assay and aligns with our objective of favoring neutral growing conditions. Accordingly, the fraction of newly accumulated alterations in essential genes – as defined by dependency maps from the Broad Institute and the Sanger Institute (*21*) (983 genes, Fig. S4B and Table S2) – was indistinguishable from that of the overall set of mutations detected in pre-MA parental PDTs (Fisher’s p-value 0.84, odds ratio 0.94).

As a control to ensure that the filtering methods applied to sequencing data did not introduce distortions in the clonal genetic divergence generated by the MA process, we used the sequencing results from both clonal PDTs and the matched parental PDT lines to derive phylogenetic trees for each tumor. The structures of the trees recapitulated the experimental design outline (Fig. S5) confirming that the observed MR heterogeneity across PDTs was driven by the diverging set of mutations accumulated during the experiment and not by analytical biases. Overall, these data indicate that different tumors (when cultured as tumoroids) accumulate *de novo* mutations at different rates and show that different clones of the same tumor are characterized by a similar MR, suggesting that MR is a unique, intrinsic and relatively stable feature of each tumor.

### Mutational signatures in *de novo* mutations

To delve into the underlying molecular processes responsible for the accumulation of new mutations in PDTs, we analyzed the mutational signatures (*22*) acquired *de novo* by each tumor during the MA experiment and compared them to those detected in the corresponding parental PDT (Fig 2A). As expected, and consistent with previous reports (*23,24*), aging-related SBS1 (arising from an endogenous mutational process of spontaneous deamination of 5-methylcytosine) was the most prevalent signature in all parental PDTs. Additionally, the mismatch repair deficiency (MMRD) SBS6 signature was enriched in the CRC0282PR MSI tumor. Interestingly, MSS tumors also displayed a signature of unclear etiology (SBS8), possibly associated with DNA late replication errors and base excision repair deficiencies (*25,26*), which has been previously reported in CRC (*24*). However, the analysis of newly accumulated mutations revealed a distinct scenario. While the MSI tumor displayed a continued presence of SBS6 (along with SBS20, another MMRD-related signature), all MSS T1 subclones showed a depletion of SBS1 and a systematic enrichment of SBS8 (Fig. 2A and Table S4). Moreover, the presence of a signature attributed to genotoxic damage induced by reactive oxygen species (*27*) (SBS18) was detected in five T1 MSS models (Fig. 2A and Table S4). This indicates that, in MSS tumors, newly accumulated mutations are predominantly sustained by mutational events that are largely independent of the SBS1-associated deamination processes typical of normal cells (*23*); instead, *de novo* acquired mutational patterns likely relate to SBS8- and SBS18-associated defects in DNA replication and repair (*25,26*). This also suggests that the prevalent SBS1 mutations detected in bulk sequencing analyses most likely arise from the set of mutations accumulated by the normal cell lineage during the period from egg fertilization to tumor initiation, rather than from events occurring later during tumor progression.

**Figure 2.**
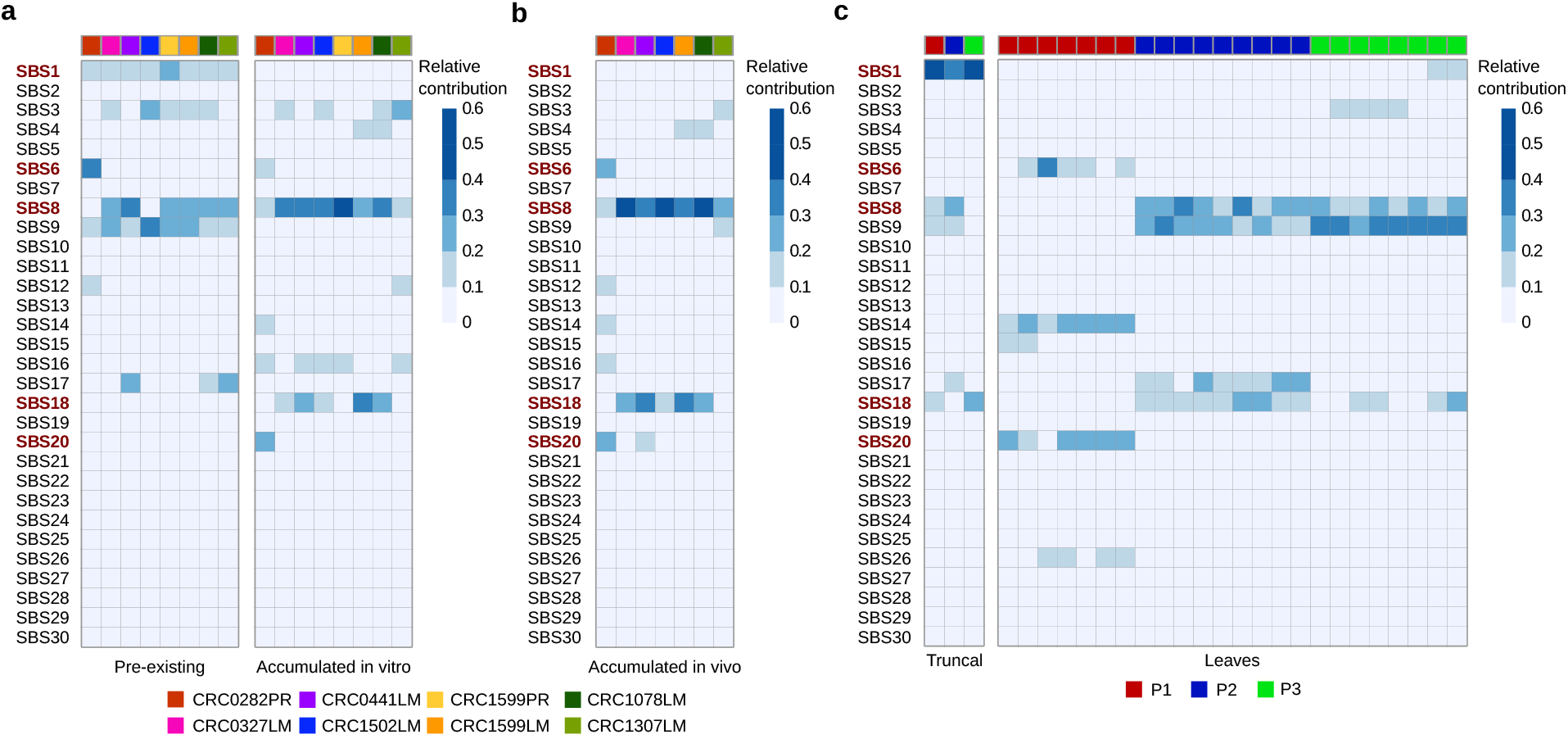
Mutational signatures contributing to mutation accumulation in CRC PDTs and PDXs. **a**, Relative contribution of COSMIC v2 signatures (SBS) to the SNVs detected in parental PDTs (Pre-existing) and to those acquired *de novo* during the MA experiment in PDTs (Accumulated in vitro). MSI-related signatures (SBS6 and SBS20) and signatures that changed during the MA experiment (SBS1, SBS8 and SBS18) are highlighted in red. **b**, Relative contribution of SBSs to the SNVs acquired *de novo* during the *in vivo* MA experiment in PDT-matched PDXs. **c**, Relative contribution of SBSs to SNVs assigned to the tumor’s most recent common ancestor (Truncal) and those private to individual single-cell clones (Leaves) in Roerink et al^20^. P1 is an MSI tumor, while P2 and P3 are MSS tumors.

To verify that the shifts in signature composition were not artefacts of *in vitro* tumoroid propagation, we set up a parallel MA experiment *in vivo* with six different PDTs; some of the cells derived from T0 clones were implanted in NOD-SCID mice and propagated as xenografts for six months; then, tumors were explanted, and T1 subclones established (Fig. 2B and Table S4). In agreement with results in PDTs, T1 subclones from xenografts showed both SBS1 depletion and SBS8/18 enrichment in MSS cases while the CRC0282PR MSI model displayed the expected SBS6 and SBS20 signatures. This consistency attests to the general relevance of these mutational changes. Again, results remained unaffected by analytical adjustments as they were confirmed by applying various signature fitting methods to both the *in vitro* and *in vivo* datasets (Fig. S6 and Table S5).

We posited that if SBS1 mutations accumulate before tumor initiation, they should be present in the tumor-initiating founder cells and persist as truncal variants, shared among different cells, in full-blown tumors. Conversely, if SBS8/18 alterations are newly generated during cancer progression, they should be detected also as branch mutations. To test this hypothesis, we took advantage of publicly available phylogenetic data of clonal tumoroids obtained from multi-regional sampling of primary CRCs (*20*). By comparing mutations that were shared among clones derived from the same MSS original tumors to those that were private to individual clones, our analysis confirmed that SBS1 mutations were predominantly found among shared mutations, while both SBS8 and SBS18 were present also among private mutations (Fig. 2C and Fig. S7A and B). We replicated these findings experimentally by generating clonal cultures from early-passage tumoroids established from three different CRC patients at the time of surgery (see Table S6 for main tumor/patient characteristics). Subsequent WGS of these tumoroids shortly after the formation of clonal colonies confirmed the enrichment of SBS1 among shared mutations, while SBS8 and, to a lesser extent, SBS18 were clearly detected also in private variants (Fig. S7C and Table S7). Not surprisingly, in both experiments SBS6 and SBS20 were ubiquitous in MSI PDTs (Fig. 2C, Fig. S7C and Table S7). Altogether, these data support the notion that SBS8 and SBS18, but not SBS1, play a major role in the accumulation of *de novo* SNVs during MSS CRC progression.

### Mutation rate stability over time

To assess the stability of MRs over longer timeframes, we performed a second MA experiment in a subset of T1 subclones from five PDTs. This involved measuring the accumulation of mutations over an additional six-month period, resulting in the collection of 25 T2 subclones that were whole-genome sequenced and compared to the corresponding parental T1 clones.

The results of the second MA experiment were consistent with the first round, both in terms of dN/dS analysis and mutational signatures (Fig. S8 and Table S8). Additionally, we found that the MR values of the T2 subclones did not differ significantly from their T1 counterparts (Fig. 3A), with the exception of the CRC0282PR MSI tumor and three T2 subclones out of 10 in one MSS tumor, CRC1502LM (Fig. 3A and Table S3). In the case of the MSI tumor, the inferred MR was already more variable across clones at T1, suggesting that the mutability of MMRD tumors is intrinsically more irregular than that of MSS CRCs. In the case of MSS CRC1502LM, the three divergent T2 subclones displayed higher mutability than the other T2 siblings and the T1 parental PDTs in terms of both MR (Fig. 3A) and absolute number of gained mutations (Fig. S9 and Table S3). Interestingly, all the divergent subclones derived from the same T1 ancestor, suggesting the presence of some inheritable traits contributing to the mutability shift.

**Figure 3.**
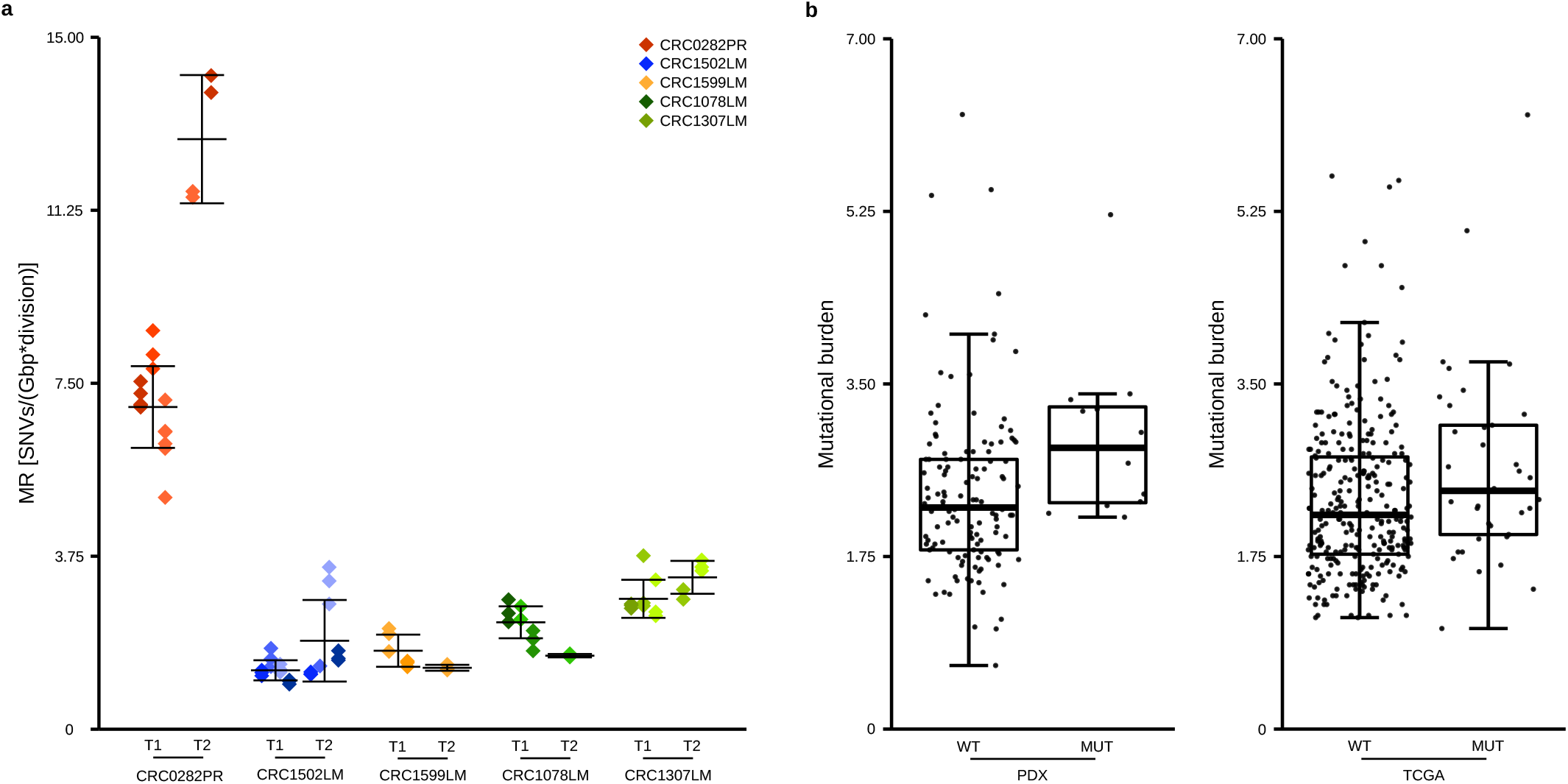
MR stability over time in MSS PDTs. **a**, Comparison between the estimate of SNV MRs computed based on the MA experiment in months 1-6 (T1) and that computed based on the MA experiment in months 7-12 (T2) in PDTs. Each dot represents the MR estimate of a clone; different shades of the same color identify subclones derived from the same parental clone. (T1, *N* = 6-15 for each PDT; total *N* = 51; T2, *N* = 3-10 for each PDT; total *N* = 25). Error bars indicate the average ±1 standard deviation. **b**, Mutational burden of tumors with or without *DNAH5* mutations in MSS samples from metastatic CRC PDXs WT, *N* = 128; MUT, *N* = 12) and in the TCGA COAD dataset (WT, *N* = 291; MUT, *N* = 37). *P* = 0.005 and *P* = 0.019 for PDXs and TCGA, respectively, one-tailed Wilcoxon rank sum test. Each dot indicates a tumor, box plots represent the overall distribution of the population, with default thresholds for whiskers (see Material and Methods for details). PDX, xenograft; MUT, mutated; WT, wild-type.

To gain insight into the potential mechanisms underlying the observed increase in MR, we concentrated on genetic variants shared among the three hypermutant T2 subclones, but not detected in the other T2 subclones of CRC1502LM or their T1 ancestors (Table S9).

Through this analysis, we identified 7 candidate mutant genes, none of which were obviously implicated in DNA repair or replication. However, a literature survey revealed that *DNAH5*, a member of the DNAH gene family of dynein motor proteins, had been previously associated with increased mutability in gastric cancer (*28*). To test whether *DNAH5* mutations correlate with a more marked mutator phenotype also in CRC, we leveraged an internally available set of whole exome sequencing (WES) data from metastatic CRC (*29*) and independently analyzed the TCGA COAD dataset (*30*). In both cohorts, *DNAH5* mutations significantly correlated with a higher mutational burden (Fig 3B and Table S9). The association was specific, as none of the other alterations common to all the CRC1502LM hypermutant clones were significantly associated with a higher mutational burden in either of the tested datasets (Table S9). Of note, the *DNAH5* mutation found in the hypermutating CRC1502LM clones (K2139E) is located in the hydrolytic ATP binding site, a genomic region conserved across vertebrates. This suggests that the resulting protein alteration may have a functional impact, although the precise mechanistic relationship with genetic instability remains unclear.

Further studies are needed to determine whether this mutation causally drives the higher MR of CRC1502LM hypermutant clones or if it is a consequence of the mutator phenotype. Nevertheless, these results strongly suggest that some kind of inheritable trait contributes to the increased mutability observed in CRC1502LM subclones.

### Accumulation of copy number alterations

By analyzing WGS data from T0, T1 and T2 clones, we found that, alongside sequence variations, copy number alterations (CNAs) also accumulated during our experiment (Fig. 4A). To quantitatively estimate the CN changes in each model, we employed MEDICC2, a tool designed to infer phylogenies based on copy number data (*31*). The distances computed by MEDICC2 rely on the number of differential CNAs among samples; therefore, the distance between a subclone and its parent (i.e., T1 versus T0 clones and T2 versus T1 clones) serves as a surrogate estimate of the number of copy number changes occurred during the MA experiment. Different from the trees obtained using SNVs, those generated by MEDICC2 only partially recapitulated the phylogenies imposed by the experimental design (Fig. S10). This discrepancy might be attributed to the smaller number of detected events, contributing to a diminished signal-to-noise ratio, as well as to the potential reversibility of CNA events. The higher variability of CNA calls was reflected also in the higher variance in the number of copy number changes accumulated by the different clones derived from the same PDT (Fig. 4B). Despite these limitations, our data revealed that there was no significant difference in the number of CNAs accumulated by the different PDTs, even when comparing the CRC0282PR MSI tumor with MSS cases (Fig. 4B). Moreover, we found no correlation between the SNV MR and the number of acquired CNAs (Fig. 4C).

**Figure 4.**
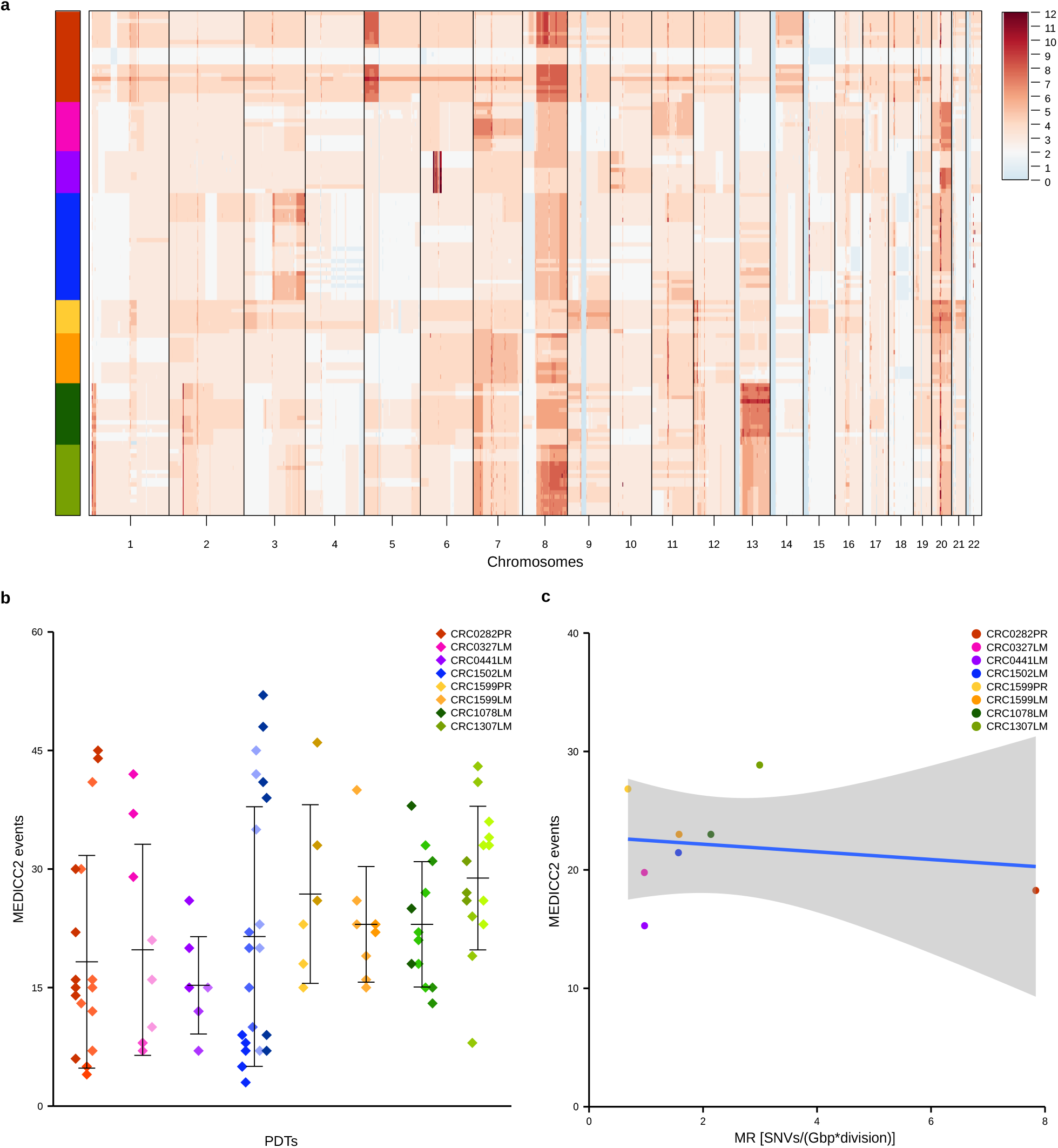
CNA accumulation in CRC PDTs. **a**, Autosomal copy number profiles of PDT clones from the MA experiment as called by Sequenza. Red and blue colors indicate gain and loss events, respectively. The profiles of individual clones (*N* = 122) are plotted as rows; the color codes on the left indicate the identity of the parental PDT (for color-coded PDT identification, see panels **b** and **c**). **b**, Estimates of de novo accumulated CNAs during the MA experiment, obtained from T1-T0 and T2-T1 distances on the phylogenetic trees. Each dot represents a T1 or T2 clone (*N* = 6-22 for each PDT; total *N* = 98); different shades of the same color identify clones derived from the same parental clone. Error bars indicate the average ±1 standard deviation. **c**, Correlation between accumulated CNAs and MRs in PDTs. Each dot represents the average estimates for an individual PDT; Spearman rho 0.06, *P =* 0.888. The shaded area represents the 95% confidence interval of the linear regression (blue line) obtained by using CNA events and average MRs as dependent and independent variables, respectively.

While CNA accumulation was assumed to take place in MSS cases, which are typically characterized by CIN, we also observed newly acquired copy number changes in the MSI case, contrary to the expectation that MSI tumors would not be affected by CIN. Previous studies have provided evidence that MSI tumors continuously acquire novel CN variants, but these variants undergo strong negative selection owing to their incompatibility with a concomitant status of DNA MMRD (*32*). The presence of accumulating CNAs in the MSI T1 and T2 subclones can be attributed to the quasi-neutral growth conditions enforced by the repetitive 100-cell bottlenecks, which were specifically designed to maintain less fit variants that might otherwise be outcompeted in a selective environment. This assumption is supported by the finding that the MSI PDTs had a trend towards a smaller number of gained CNAs, indicative of lower CIN tolerance (Fig. 4B). Interestingly, the clones of the TP53 wild-type model (CRC0441LM) also tended to acquire fewer CNAs, likely due to barriers against karyotype imbalance in the context of a functional P53 pathway.

### Higher mutation rates in metastatic tumors

Building on the observation that MR appears to be a unique and inheritable feature of each cancer cell, we asked whether MR could be subject to selective pressure during cancer progression. Upon examining pairs of PDTs originating from the same patient (Fig. 1B and Table S3), we noticed that the lesion arising later during cancer progression exhibited a higher MR. Specifically, the MR of a liver metastasis (CRC1599LM) was 2.3-fold higher than that of the corresponding primary tumor (CRC1599PR). Similarly, the MR of CRC1307 (a metastatic recurrence that became clinically evident approximately one year after removal of a previous liver metastatic nodule) was 1.4-fold higher than that of CRC1078 (the earlier metastasis from the same patient) (Table S3). To extend our analyses beyond the cohort of PDTs used for MA experiments, we decided to evaluate the distribution of subclonal VAFs in a larger set of tumors, with the underlying rationale that the steeper the slope of the cumulative distribution of subclonal neutral VAFs, the higher the MR. In accordance with this notion, the number of subclonal SNVs in clones of CRC1599PR, CRC1599LM, CRC1078LM and CRC1307LM significantly correlated with the MR computed by means of MA experiments in the same models (Fig. 5A and Table S10). This correlation was consistent when analyzing WES data of the PDXs from which CRC1599 PDTs were derived; the PDX derived from the metastatic lesion displayed a steeper distribution of subclonal variants than the PDX derived from the primary tumor (Fig. 5B). On this ground, we employed a heuristic algorithm that provides a proxy of MR by fitting a linear regression model to the cumulative distribution of subclonal variants from bulk tumor sequencing data (*14*). We applied this method to a dataset comprising 27 pairs of high-depth (300x) WES results obtained from 54 PDXs derived from primary CRCs resected synchronously with their corresponding metastases. Among these, 16 pairs met the quality criteria required for the analysis (R^2^ > 0.9 and > 10 subclonal mutations in both samples) when setting standard VAF thresholds (0.12 < VAF < 0.24) to define subclonal variants. With these parameters, we obtained a median slope of the linear regression (an indirect indicator of the MR per effective cell division) that was significantly steeper in the metastases than in their matched primary tumors (11.8 and 16.6; *P* = 0.00015 by paired Wilcoxon signed rank test), with 87.5% of the cases displaying increased MR in the metastatic deposit (Fig 5C and Table S11). This outcome remained consistent when using a wide range of VAF thresholds to define subclonal mutations (Fig. S11A and Table S11). Moreover, the same result was obtained using the raw count of subclonal mutations as a surrogate of MR (Fig. S11B) and remained significant even after the exclusion of the two PDX pairs showing the largest difference (Fig. S11C). Notably, the primary-metastatic tumor pair in which the MR was measured experimentally and all pairs in which MRs were inferred computationally originated from synchronously resected samples, thereby eliminating the possibility that the higher MR detected in metastases was influenced by the mutagenic effects of intervening chemotherapy. Overall, these data suggest that a higher MR is selected during CRC metastatic progression.

**Figure 5.**
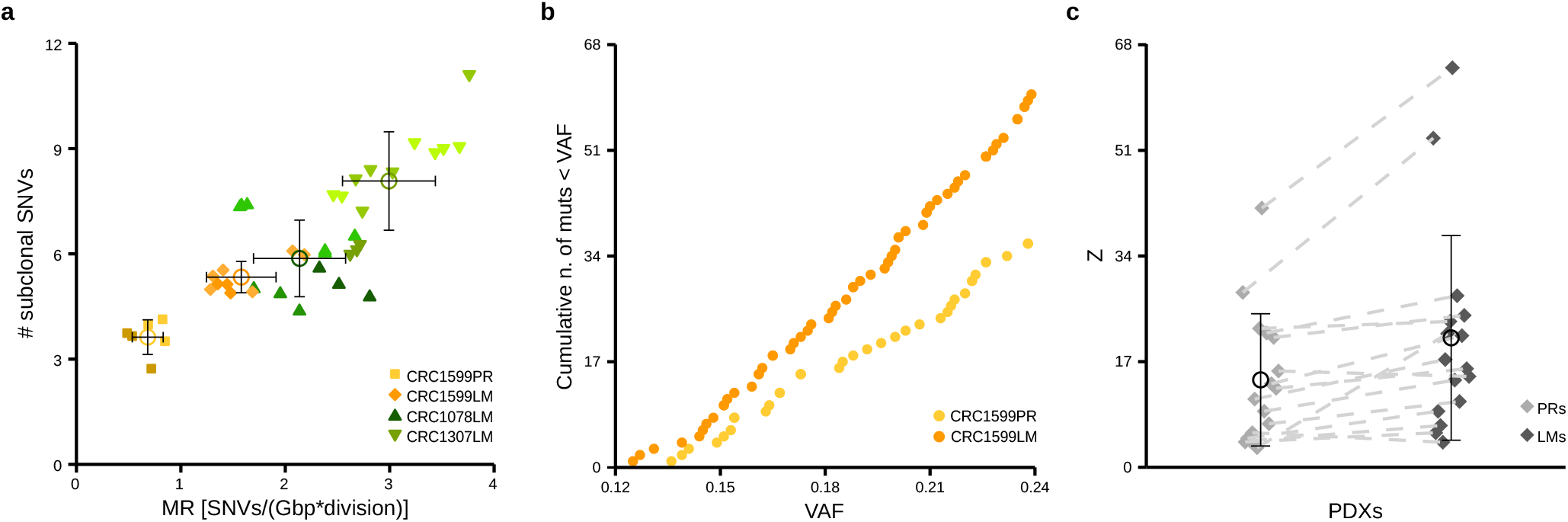
MRs in primary tumors and paired metastatic lesions. **a**, Correlation between MRs and the total number of subclonal SNVs in 41 clones from 2 pairs of PDTs, each consisting of less and more advanced tumors from the same patient. Each dot represents a clone; different shades of the same color identify T1 and T2 subclones derived from the same T0 clone. Error bars indicate the average ±1 standard deviation for each PDT; empty circles indicate the mean. CRC1307LM (*N* = 14) vs CRC1078LM (*N* = 12), *P* = 4.038e-05; CRC1599LM (*N* = 9) vs CRC1599PR (*N* = 6), *P* = 0.0004, Wilcoxon rank sum test. **b**, Cumulative distribution of subclonal mutations in PDXs from a primary tumor (CRC1599PR) and its matched synchronous metastasis (CRC1599LM). Each dot represents the number of subclonal mutations with an observed frequency < VAF. **c**, Numerical values of the slopes (Z) for the linear regression between cumulative frequency distribution and inverse frequency of subclonal mutations (0.12 < VAF < 0.24) in PDXs derived from matched synchronous resections of primary and metastatic tumors (*N* = 16). Dashed lines connect primary tumors (PRs) to their corresponding liver metastases (LMs). *P* = 0.00015, one-tailed paired Wilcoxon signed rank test. Error bars indicate the average ±1 standard deviation; the average is represented by an empty circle.

## Discussion

This study provides a quantitative assessment of MRs in a cohort of eight different PDTs illustrating prevalent molecular subtypes in CRC. To our knowledge, this represents the first quantitative comparative measurement of MRs using a controlled experimental design across multiple cancer models.

The reported MR estimates from bulk WGS data of MSS CRC tumor samples fall within the range of 10-31·10^-9^ mut/(bp gen) (*10,11*). In contrast, the MR values derived from our MA assays are around one order of magnitude lower, with an overall average of 1.6·10^-9^ mut/(bp gen) in the MSS models considered. While DNA sequencing analyses have to rely on mutational distance distributions to infer cell survival and death rates per division (*11*), we used empirical measurements to enumerate DNA replication and death rates over experimental time. We surmise that computational approximation based on cell net growth, without explicit information on cellular death rates, may not fully account for the number of unsuccessful cell divisions, which nonetheless contribute to the number of DNA replications separating two mutations. Another possibility is that, when dealing with bulk sequencing data, the concomitant expansion of multiple clonal lineages within patients’ tumors could favor an overestimation of the number of subclonal mutations attributed to the neutral expansion of the founder cell, again leading to higher MR estimates.

The deployment of patient-derived clonal tumoroids for MA assays is unprecedented and is expected to yield results that more faithfully reflect the clinical setting of spontaneous tumors in patients. Our work shows that MRs were heterogeneous among the different models examined and, for the most part, similar between independent clonal progenies derived from individual cells of the same model. Therefore, MR appears to be an inherent and evolvable characteristic of each tumor (*33*). Experimental evidence also suggests that MR can be dictated by inheritable traits, as exemplified by the *de novo* acquisition and fixation of a potentially functional mutation in the *DNAH5* gene in subclones derived from the same ancestor and displaying higher mutability than the parental tumoroid in a CIN model. This correlation is strengthened by the observation that, in two clinical datasets, *DNAH5* alterations were enriched in CIN CRC tumors with a higher mutational burden. The notion that the gain of hitherto unappreciated heritable variants may propel the accumulation of new mutations also in tumors that do not harbor well-established drivers of DNA hypermutability (such as mutant forms of mismatch repair enzymes and DNA polymerases) is in line with previous reports hinting to high MRs as a general occurrence in tumors compared with normal tissues (*34-36*).

In our hands, normal colonocytes did not survive the repeated single-cell dissociation and cloning steps imposed by the MA protocols, preventing a direct comparison of the MR of cancer cells with that of normal cells. However, *in silico* and experimental MR estimates in lower organisms and normal human tissues range between 0.05 and 0.46 *10^-9^ mut/(bp gen) (*11,37*), well below the MR values calculated in our MA assays. Moreover, the assumption that CRCs are endowed with higher MRs than normal colonic cells is consistent with the emergence of the SBS8 and SBS18 mutational signatures, as revealed by our MA experiments. These signatures, which are both associated with genotoxic damage and defects in DNA replication and repair, were apparent in newly acquired mutations, but they are barely detectable in normal cells (*38*). Thus, ongoing mutational processes related to DNA mutability appear to be a peculiarity of CRCs not shared with normal tissues.

Conversely, while the SBS1 aging signature was clearly evident in parental PDTs, its contribution to accumulated mutations was negligible, even after one year of continued growth. This finding suggests that the number of SBS1 mutations found in the tumor bulk may largely be traced back to the normal cell that originally initiated the tumor, rather than post-transformation mutational processes. From this perspective, the number of SBS1 mutations could be interpreted more as a proxy of the age of the patient at the time of tumor initiation than as a measure of the age of the tumor. This insight may have relevant ramifications for estimating the dynamics of cancer progression and their relationship with clinical outcome.

Experimental findings from MA assays and computational analysis of VAF-based MR estimators documented higher MRs in metastatic lesions than in matched primary counterparts, arguing for positive selection of cells with progressively increasing mutability during cancer dissemination. If confirmed in larger patient cohorts, this observation may impact the way drug response and resistance are modelled biologically and addressed clinically. For example, if cells composing primary tumors or early subclinical micrometastases mutate slower than cells of established metastases, the time to resistance to targeted drugs (for the same number of cell individuals) should be longer for “younger” lesions. Additionally, different MRs may implicate different rates of neoantigen generation, which would make tumor stage a potentially important parameter for predicting the efficacy of immunotherapeutic approaches.

In conclusion, our study highlights the relevance of DNA sequence mutability as a fundamental biological property that typifies the identity and evolutionary strategies of CRC tumors, including CIN tumors without obvious drivers of mutational instability, and introduces new metrics that may contribute to a better understanding of the principles underlying tumor diversification over time.

## Supporting information

Methods, Supplementary Figures and list of Supplementary Tables

## Acknowledgements

We thank Mauro Papotti, Gianluca Paraluppi, Alessandro Ferrero and Serena Perotti for sample acquisition; Massenzio Fornasier and Arianna Russo for veterinary assistance; Fabrizio Maina for animal husbandry; Raffaella Albano, Stefania Giove, Lara Fontani, and Laura Palmas for technical assistance; and Daniela Gramaglia and Mauro Paschetta for secretarial assistance.

## Author contributions

Conceptualization was the responsibility of A.Be. Study design was carried out by A.Be., L.T., E.G., V.V., I.C., A.S., M.C.L, G.U., A. Ba.

V.V performed all the tumoroid culturing and material extraction with help from I.C., B.L and F.G. F.S. acquired and quantified EdU stainings. V.V., E.R.Z. and F.C. performed *in vivo* experiments. Analysis of WGS data and MA pipeline development was the responsibility of E.G., with support from A.Be., A.S., G.C., B.C., G.U., S.B., M.V. and S.P. G.C. performed the copy number analysis with haplotype phasing and phylogenetic reconstructions. E.G., S.B. and M.F. performed visualization. M.V. carried out data deposition. Funding acquisition was undertaken by A.Be. and L.T. A.Be. and L.T. supervised the work. Writing of the original draft was done by E.G., M.C.L., S.P., G.C., V.V, A.Be and L.T. Writing review and editing were performed by all authors.

## Data and materials availability

GATK best practice pipeline, Sequenza and Platypus multi-sample calling procedure were implemented with Snakemake and available at https://github.com/vodkatad/snakegatk, with Docker images available for different rules. The Snakemake workflow to identify gained mutations, compute MRs, perform signature analyses is available at https://github.com/vodkatad/AF_spectra. Specifically all the panels of the main Figures and the majority of Supplementary Tables and Figures are reproducible with specific Snakemake rules whose inputs have been uploaded on https://github.com/vodkatad/AF_spectra (release M1, directory dataset_Figures_Tables). The whole pipeline leading to those inputs can be re-run downloading raw data from EGA (accession codes will be available before publication, if required raw data can be made available during revision); intermediate, but privacy restricted files, such as vcfs, can be requested to the authors via e-mail.

