## Supplementary material for "Heterogeneity and evolution of DNA mutation rates in microsatellite stable colorectal cancer": Methods, Supplementary Figures and list of Supplementary Tables

#### Materials and Methods

##### Patient-derived tumoroid cultures

CRC tumoroids were established from PDXs described in [39]. Tumor specimens (~50mm$^3$) were chopped with a scalpel and washed with PBS. After centrifugation, the final cell preparation was embedded in Matrigel® (Corning) or Cultrex Basement Membrane Extract (BME Type II or Ultimatrix RGF BME, R&D Systems) and dispensed onto 24-well plates (Corning). The cells were incubated for 10-20 minutes at 37°C, before adding culture medium as follows: Dulbecco's modified Eagle medium/F12 supplemented with penicillin-streptomycin, 2 mM L-glutamine, 1mM n-Acetyl Cysteine, B27 (Thermo-Fisher Scientific), N2 (Thermo-Fisher Scientific) and 20 ng/ml EGF (Sigma-Aldrich). Tumoroids were maintained at 37°C in a humidified atmosphere of 5% $CO_2$ and regularly tested for Mycoplasma infections. In order to ensure matching sample identity with the original human specimen from which they originated, all the models were genotyped by means of the MassARRAY Analyzer 4 (SEQUENOM® Inc, California) using a 24 SNPs custom genotyping Panel (Diatech Pharmacogenetics). Culture expansion and biobanking materials were managed using the Laboratory Assistant Suite[40].

Single-cell cloning was performed by seeding disaggregated tumoroids at limiting dilution in 96-multiwell plates, followed by microscopic inspection to select wells containing individual single-cell clones. Shortly after derivation (approximately 6 weeks), some cells from each clone were archived under viable conditions and others underwent DNA extraction for sequencing. The remaining cells were further propagated to allow for the accumulation of *de novo* mutations. We periodically induced bottlenecks by dissociating clones and replating them at a density of approximately 100 random individual cells every two weeks to favor neutral evolution. Over the course of the experiment, cells were randomly sampled to undergo quantification of DNA replication rates through EdU staining.

To obtain clonal populations from tumors at the time of surgery, tumoroids established from fresh resections were processed shortly after collection to obtain single-cell clones, as described above.

##### DNA extraction

Total DNA was extracted using the Maxwell® Instrument (Promega) following manufacturer's instructions. Specifically, 30 µl of Proteinase K (PK) Solution, 270 µl of PBS and 300 µl of Lysis Buffer were added to the pellet of tumoroids, then each sample was vortexed 10'' seconds and incubated at 56°C for 20 minutes before proceeding with the extraction.

##### EDU staining

Cell proliferation assays were performed using a Click-iT Plus EdU Imaging kit (Thermo Fisher) according to the manufacturer's instructions. Subconfluent cells were supplemented with 5'-ethynyl-2'-deoxyuridine (EdU) at a 10 µM final concentration for 3 hours. After that, cells were harvested, trypsinized and dissociated at single-cell before collection in PBS (1ml). The cell suspension was centrifuged for 5 min (1200 rpm) and then fixed with 1 ml 4% PAF for 10 min at 4°C. Following fixation, cells were washed and then permeabilized with 0,5% Triton X-100 in PBS

(1ml) for 20 min at room temperature. The permeabilized cells were washed two times with 3% BSA in PBS (1ml). Cells were then incubated with 500 µl Click-iT reaction buffer for 30 min at room temperature, protected from light. Following the Click-iT reaction, cells were rinsed twice with 1 ml of 3% BSA in PBS and then incubated with DAPI (4',6-diamidino-2-phenylindole) 1:1000 in PBS for 5 min at room temperature. EdU-stained cells were then washed again with PBS and mounted for imaging.
Images were captured using a Nikon eclipse microscope and analyzed with 'ImageJ' software, the percentage of EdU positive cells were considered reliable when at least 500 nuclei were scored.

**In vivo xenografts passaging**

Tumor implantation and expansion in NOD-SCID mice was performed as previously described[41]. Tumor size was evaluated once-weekly by caliper measurements and the approximate volume of the mass was calculated using the formula $4/3\pi \cdot (d/2)^2 \cdot D/2$, where d is the minor tumor axis and D is the major tumor axis. When the tumor larger diameter exceeded 1.5 cm, tumors were explanted, fragmented, and passaged in new mice. After six months of continued propagation, xenografts were explanted and dissociated mechanically using the GentleMax Dissociator (Miltenyi Biotec), followed by enzymatic digestion of the extracellular matrix with the Human Tumor Dissociation Kit (Miltenyi Biotec), according to the manufacturer's protocol. Cell suspensions were then subjected to single cell cloning as described above.
All animal procedures were approved by the Ethical Commission of the Candiolo Cancer Institute, IRCCS – FPO and by the Italian Ministry of Health (Authorization # 37/2022-PR).

**Tumoroids birth rates estimates**

The estimate of the number of DNA replications per day was calculated as 8 times the fraction of EdU positive cells, which represent the ones that were in the S phase of the cell cycle during the 3-hour incubation period. To calculate the total number of DNA replications during the MA experiments, for each T0 clone the average number of replications per day was obtained and then multiplied by the total number of days of mutation accumulation. The resulting number of DNA replications was used to calculate the MR (*gens*, see below). We also estimated the effective division rates by an exponential fitting of the cell counts (*c*) obtained before each 100-cells bottleneck. The ratios between $log_2\left(\frac{c}{100}\right)$ results and the number of days between successive bottlenecks were averaged to obtain, for each model, the number of population doublings per day. As expected, the resulting number of total generations for each clone was smaller than the one obtained via EdU based measurements of cell divisions, since population-doubling estimates capture effective growth, which is also affected by cell deaths.
For one clone (CRC1078LM-02) for which EdU estimates were not available, we resorted to averaging the number of divisions obtained for the other two T0 clones originated from the same PDT.

**Specimen collection and annotation**

Tumor samples were obtained from patients treated by liver metastasectomy at the Candiolo Cancer Institute (Candiolo, Torino, Italy), Mauriziano Umberto I (Torino, Italy) and San Giovanni Battista - Molinette (Torino, Italy). All patients provided informed consent, samples were procured and the study was conducted under the approval of the review boards of the institutions. Clinical and pathologic data were entered and maintained in our prospective database.

##### WGS sequencing and MA clones identifiers

Genomic DNA was sequenced on Illumina NovaSeq or HighSeq by Biodiversa SRL with a standard protocol for WGS library preparation using PCR amplification. We aimed at a mean depth of 30x with 150bp paired reads (90 Gbp per sample). The list of all the sequenced clones and bulk samples, alongside their matched normals, is available in Extended Data Table 11. Every sample id begins with a 7 letter alphanumeric code identifying the surgical material from which the original xenograft was derived, followed by two letters – PR (primary tumor) or LM (liver metastasis) – to distinguish the synchronous lesions coming from the same patient (i.e. CRC1599PR and CRC1599LM) and their corresponding germline tissues – NM (normal intestinal mucosa adjacent to the primary tumor) or NL(normal liver) The letter in position 10 indicates whether the DNA was derived from a tumoroid (O, organoid), a tumor grown in immunocompromised mice (X, xenograft), or if it was directly extracted from the surgical material (H, human).

For clones that underwent MA experiments, their phylogenetic relationship is coded by hyphen-separated descriptors (positions 11-18) as follows:

- For T0 clones, XX-0; where XX indicates the identity of the clone and 0 indicates T0.
- For T1 subclones, XX-1-Y; where XX indicates the identity of the parent T0 clone, 1 indicates T1 and Y indicates the identity of the subclone.
- For T2 subclones, XXX-2-Y; where XXX indicates the identity of the parent T1 clone, 2 indicates T2 and Y indicates the identity of the subclone.
- For clones that underwent MA in vivo, XX-MY-Z; where XX indicates the identity of the parent T0 clone, MY indicates the in vivo propagation method (MA, transplantation of peripheral tumor fragments from mouse to mouse; MC, transplantation of central tumor fragments from mouse to mouse; MI, transplantation of well-mixed tumor cell suspensions from mouse to mouse).

Whenever possible, we generated three T0 clones and nine T1 clones (three for each T0). Not all clones survived the whole experiment therefore there are exceptions to this rule (e.g. CRC0441-10-1-C is the only T1 clone available for the ancestor CRC0441-10-0). For six models, 1-3 T0 clones were also propagated in vivo and 3-9 subclones were analyzed at T1. For five models, 6-15 T1 subclones were propagated for further six months and 3-10 T2 subclones for each T1 were subjected to WGS to monitor MR stability over time.

##### WGS SNV and small indels single-sample calling

We followed the GATK Best Practices for somatic mutation calling with Mutect2[42–44], performing the standard steps of alignment (bwa[45] version 0.7.17-r1188), marking of duplicates (picard 2.18.15), quality recalibration (GATK 4.1.4.0) and mutation calling against matched normal. For parental (pre-cloning) samples we used all the default parameters, while for the clones deriving from a single cell we used ''--default-af 0" to avoid the automatic categorization as germline variants of all the mutations with an allelic frequency of 0.5.

As a reference genome we used GRCh38, including some additional known contaminants (eg. HPV), made available by the GDC portal[46].
We used different resources, as suggested by the Best Practices[47], to refine the calling procedure: a panel of normals, based on 1000G, was used to filter out sequencing artefacts; the NCBI dbSNP polymorphisms was used for quality recalibration and finally we retained only mutations located in the list of mappable regions, to avoid mapping biases in centromeres and highly duplicated regions.
We used mosdepth[48] (version 0.2.3) to segment the genomes based on the depth of sequencing.
Basic QC was performed using FastQC[49] (version 0.11.7), picard CollectWGSMetrics and samtools[50] (version 1.9) - mean, sd and median coverage are listed in Extended Data Table 12. All the samples provided a mean coverage of 19x or more, which met the requirements for further analyses. Two samples (CRC0282-01A-2-3 and CRC0282-07E-2-2) were excluded from further analyses due to a potential cross contamination (> 0.001) based on GATK CalculateContamination tool (see Extended Data Table 12).
The mutational calling pipeline was all run on the University of Turin HPC cluster OCCAM[51] via snakemake[52] (version 5.4.0) and docker. Mutations were functionally annotated with annovar[53] (version 2018-04-16).

**WGS CNV calling**
We used Sequenza[54] (version 3.0.0) to analyze the alignments of each tumor and its matched normal, using the marked duplicates obtained for the mutation calling pipelines as an input. We binned the genome in 200bp windows and used all the default parameters, based on which the SNPs represented by dependable base calls (quality > 20) were exploited to compute the B allele frequency. We verified that, as expected in case of pure cancer-cell cultures, the optimization procedure ended up with solutions implying ~100% cancer cell purity.
For 2 clones of CRC0327 (02-1-E, 02-1-I) Sequenza inferred a wrong ploidy (6), which resulted to be triploid following manual curation of the normalized sequencing depth evaluation of the B-allele frequency distribution. For these clones we forced the sequenza fit algorithm to assume 100% cancer cell purity and ploidy = 3.
Sequenza CN calls were adopted for MR computations (see below) as we opted for a more coarse-grained segmentation.

**WGS CNV calling using haplotype phasing and phylogenetic analysis**
To increase our sensitivity to subclonal events we also performed CNV calling using haplotype phasing and used the results to perform phylogenetic analysis based on CNVs.

<u>Phasing references</u>
Reference vcf files were downloaded from the 1000 genomes European Bioinformatics Institute (EBI) resource[55] (ALL genotypes, dated 2017-05-03, release 2013-05-02), mapped to GRCh38. We took only female X chromosome information to ensure the reference was biallelic. Positions with unknown genotypes (e.g., ./0, 0/., ./.) and duplicate entries were removed, as were entries including the “END=” label in INFO. Lastly, mappability was calculated using GEM library[56] and only uniquely mapping variants were used.

Performing phasing

Remaining variants were used to genotype the normal BAM files of each patient using Platypus[57] (genotyping mode, --getVariantsFromBAMs=0). Only PASS variants were then used. Next Beagle[58] was used to haplotype phase the germline single nucleotide polymorphisms (SNPs) of the patient (impute = false, burnin = 5000, iterations = 10000) using plink maps provided by Beagle for GRCh38.

Copy number calling using haplotype predictions

Allele frequency and depth ratios were obtained using the SEQZ output from Sequenza[54]. GC content was normalized using the *gc.norm()* function in Sequenza. Only heterozygous SNPs with a minimum depth of 25 reads in the normal sample were used for analysis. $Log_2$ of the depth ratio was calculated and the median $log_2$ depth ratio value subtracted from all SNPs to normalize to the median.

To call CNAs using Beagle haplotype phasing, we utilized an adapted version of the Battenberg[59] workflow. Patient-specific heterozygous SNPs were subset for overlap with the phasing reference SNPs. B-allele frequencies (BAF) were then calculated according to the haplotype phasing such that the BAF represented a single haplotype prediction. A single chromosome prediction set of SNP allele frequencies was segmented using piecewise-constant fitting (*selectFastPcf*) from the Battenberg package (KMIN = 1, gamma = 3 * *sdev*, yest = T). The *sdev* was calculated using the *getMad()* function (k=25) on the mirrored BAF per chromosome (min. *sdev* 0.002). The BAF of SNPs belonging to segments with a mean BAF less than 0.5 were subtracted from 1 to fully phase the BAF after haplotype block detection. We then initially used joint segmentation to segment the log2ratio and the phased BAF, to detect changes in total copy number and allele balance simultaneously, using the multiple sample piecewise constant fitting function in the copy number package[60] (*multipcf*, gamma = 10). We then removed SNPs in segments that had mean values less than 0.45, indicating poor phasing, and repeated the joint segmentation using stricter settings (gamma = 20). We then normalized the log2ratio values using the *normalize()* function in CGHcall[61] (method = “median”, smoothOutlier = T). Copy numbers were then fit using the *fit.copy.number()* function in ASCAT[62] searching for high tumor purity fits only (dist_choice = 0, ascat_dist_choice = 2, min.ploidy = 1.6, max.ploidy = 5.5, min.rho = 0.95, min.goodness = 0.63, uninformative_BAF_threshold = 0.51). Subclonal copy number cancer cell fractions were determined using the *callSubclones()* function in Battenberg (segmentation.gamma=NA, siglevel = 0.05, maxdist = 0.01, noperms = 1000, seed = 1, calc_seg_baf_options = 1).

Ploidy search ranges were reduced in CRC0282-01-0, CRC0282-07-0, CRC0282-07-1-A, CRC0282-07-1-D and CRC0282-07-1-E after manual assessment of the presence of tetraploidy (min.ploidy = 3, max.ploidy = 5) and purity and ploidy was pre-set entirely for CRC0282-01-0 and CRC0282-07-1-D (preset_rho = 1, preset_psi = 3.6).

Phylogenetic analysis using MEDICC2

Only *in vitro* samples were used for phylogenetic tree analysis. To reduce the resolution of the calls to ensure phylogenetic trees were not dominated by small changes, we binned the segments into 1 Mbp bins across the genome, taking the most prevalent CNA status per segment as determined by Battenberg. In the event of multiple segments overlapping with a bin, we took the median copy number weighted by the overlap of the segment with the bin. Bins that had a copy number of 1000 were removed as were bins without any overlapping segments in a minimum of one of the samples. Phylogenetic trees were then calculated using MEDICC2[31].

**Multi-sample mutational calling and MRs computation**

To reliably identify the mutations that accumulated during the experiment we considered only mutations in regions that are covered with at least 1 read with mapping quality > 20 in all the clones derived from the same model (i.e. all the clones at the beginning and at the end of the 6 months accumulation period).
We collected all the SNVs identified by Mutect2 in at least one of the clones and then used Platypus[57] (version 0.8.1.2) in Combined Genotyping/Calling Mode mode (subcommand callVariants with options "--minPosterior=0 --getVariantsFromBAMs=0 --minReads 1") to score positive for the mutation all the clones in which at least one supporting read was detected. When comparing the T1 (end of six months) and T0 (beginning) clones we focused only on mutations located in regions with CN comprised between 1 and 3, to avoid sensitivity biases.

Therefore, a mutation was defined as 'accumulated' if:

- it was detected by Mutect2 in any of the clones
- at least 1 supporting read was found in a specific subclone at the end of the experiment
- no supporting reads were found in the originating clone at the beginning of the experiment

The mutation rate for a T1 (or T2) clone is then defined as:

$$\frac{Ng}{(gens \cdot L)}$$

Where *Ng* is the number of accumulated mutations, *gens* is the number of DNA replications that took place during the experiment (estimated by means of EdU incorporation, as previously described), and *L* is the length of the considered genome, corrected for copy number status.
All the region overlaps were computed with bedtools[63] (2.27.1).

**Alternative approaches for MR computation**

We chose a very lenient filter on coverage to obtain an extensive list of accumulated mutations, to ease their subsequent characterization. As a sanity check, to confirm that this choice did not introduce biases in our MR estimates, for two PDTs we computed the MR also using a more stringent threshold. Specifically, we restricted the analysis to regions with a coverage equal or higher than 20x in all the clones. In these conditions, we detected roughly half of the mutations identified when adopting less stringent parameters. However, we obtained very similar results in terms of MR (fold changes between average MRs for each PDTs: 1.07 and 1.18 and Spearman correlations between clones' MRs for the two PDTs: 0.98, see Extended Data Fig. 12A).
Similarly, we extended the mutation calls to clones in which a single mutated read was detected, when the same mutation had been called by Mutect2 in a related clone. The main line of reasoning behind this second lax choice on multi-sample mutation calling was in line with what is known about calling mutations in related samples[57] and to limit the biases due to potential false negatives caused by the limited depth of our sequencing.

To test whether these choices could result in any biases related to the overestimation of the accumulated mutations (e.g. the inclusion of subclonal variants or sequencing artefacts as accumulated mutations) we also applied a more conservative approach to define the set of accumulated mutations. Specifically, we aimed at selecting only the variants whose VAF and CN status were compatible with being clonal at the end of the MA experiment. To achieve this, we used single-sample Mutect2 calls from T1 clones and we retained only those with a number of supporting reads and total coverage that were compatible with the expected VAF for clonal variants. The expected VAF and its standard deviation ($\sigma$) were defined using the binomial distribution with probability of success ($s$) determined by the copy number status: i.e.:

$$s=\frac{1}{CN}$$

with sd:

$$\sigma=\sqrt{s\cdot(1-s)}\cdot\sqrt{n}$$

where $n$ is the total coverage and $CN$ the copy number status of a given SNV.
Only SNVs whose altered reads were compatible with the average for this binomial distribution $\pm 1\sigma$ were considered:

$$s\cdot n-\sigma<alt_{reads}<s\cdot n+\sigma$$

$$s-\frac{\sigma}{n}<\frac{alt_{reads}}{n}<s+\frac{\sigma}{n}$$

$$s-\frac{\sqrt{s\cdot(1-s)}}{\sqrt{n}}<\frac{alt_{reads}}{n}<s+\frac{\sqrt{s\cdot(1-s)}}{\sqrt{n}}$$

where $alt_{reads}$ are the total reads supporting the SNV.
Therefore, a mutation was counted as “accumulated” only if the above conditions were satisfied and it was not detected in the originating clone at the beginning of the experiment. The same limits in CN and coverage as our original MR calculations, were applied.
As expected the MRs obtained were systematically lower than those presented in the main text; however, inter-tumor differences were overall comparable (Extended Data Fig 12b) and the two distributions of average MRs (for each PDT) correlated well (Spearman correlation 0.9, see Extended Data Fig 12c). In particular the differences in MRs between the synchronous primitive and metastatic lesions and the CRC1502LM hypermutant T2 clones were consistent independent of the calling approach selected.

##### dN/dS and regional enrichments

For dN/dS analyses we used dNdScv[64] (version 0.0.1.0). All the mutations accumulated by the different clones originated from the same PDT were aggregated to obtain more robust estimates.
Annotatr[65] (version 1.12.1) was used to define the genomic localization of accumulated and truncal mutations.

##### Phylogenetic reconstruction

SNVs and indels identified by means of the multi-sample procedure (for clones) and by the standard Mutect2 pipeline with VAF > 0.1 (for the parental tumoroids) were used to compile a digitalized matrix. Phangorn[66] (version 2.6.3) was then used to derive a phylogeny from the matrix, choosing the best tree (out of 10000 bootstraps) with the parsimony ratchet method, the root was fixed at the matched normal sample.

##### Signatures analyses

Mutational signatures fitting was performed on the aggregated list of all the mutations accumulated by clones originated from the same PDT in a MA experiment using the MutationalPattern package[67] (version 1.12.0) and the list of Cosmic Single Base Substitution Signatures (v2). The contribution of the fit for each signature was represented as the fraction of the total signal, and cosine similarity between the original mutational profile and the one obtained by the fitting procedure was used to evaluate the quality of the fit.
We used Signal[24] (signature.tools.lib version 2.4.1, R zversion 4.3.0) and sigfit[68] (version 2.2, R version 4.3.0) as alternative methods to determine signatures exposures. Moreover, to evaluate the robustness of our results under different analytical scenarios, we chose a different set of reference signatures for Signal, namely the ones detected in Degasperi et al. in the GEL colorectal cohort.

##### TCGA and WES mutational burdens analyses

The total number of mutations for each sample in the TCGA dataset together with the list of genes with coding (missense/nonsense/frameshifts) somatic alterations for the same samples were downloaded from cbioportal (TCGA, PanCancer Atlas 2018[69]). The total mutational burden was calculated by dividing the total number of mutations by the length of the most common WES capture kit adopted by TCGA for COAD/READ (SeqCap EZ HGSC VCRome, S1 Table in[70]). The analysis was performed only on CIN samples to avoid bias due to hypermutated tumors. The same procedure was applied to a previously published dataset of PDXs from our lab[29].

##### Identification of private and shared SNVs for single cell derived PDTs

We adopted the same multi-sample mutational calling procedure described above for clones derived from a single tumor (Extended Data Table 14), to identify private and shared SNVs. Phylogenetic reconstructions on SNVs identified by Roerink et al.[20] were used to distinguish private and shared SNVs in their clones - we defined as 'Leaves' all SNVs assigned to leaf nodes in Extended Data Figure 3 / Supplementary Data S4; while we labeled as 'Truncal' the ones assigned to the MRCA node of all the tumor clones (nodes '17' and ‘17_not_timed_to_WGD', '17_postWGD', '17_preWGD').

**WES sequencing on primitive-metastatic pairs and MR inferences from bulk sequencing data**

WES for 54 matched Primitive-Metastatic xenografts was performed with the xGen Exome Hyb Panel v2 capture kit by IDT and sequenced on Illumina NovaSeq or HighSeq by Biodiversa SRL.

We aimed at a mean depth of 300x with 150bp paired reads; mean, sd and median coverage are listed in Extended Data Table 15. Mouse-derived reads were filtered using Xenome[71] (version 1.0.0) with default parameters and k-mer indexes obtained from GRCh38 and mm10. Mutation calling was performed using the same pipeline and software described for WGS, specifying the IDT targeted exons and a padding around them of 100bp. The fitting procedure for the cumulative distribution of subclonal variants described in Williams et al.[14] has been re-implemented in R.

**Statistical analyses**

The number of available data-points (e.g. different clones) is reported in the figure legends, alongside the adopted statistical tests and metrics. Boxplots extend from the first to the third quartile with the median shown as a horizontal line, whiskers extend to the more extreme point within |1.5*IQR| from the box borders; all individual points are shown. All graphs and statistical analyses were obtained with R (version 3.6.3, unless specified otherwise), its base packages and the following libraries: ggplot2[72] (version 3.3.0), ggpubr (version 0.2.5) and pheatmap (version 1.0.12.), with the exception of the CN heatmap (Fig 4A) that was made with python3/matplotlib (versions 3.7.3 and 3.4.3). Individual panels were assembled and subjected to cosmetic fixes using Inkscape.

a b

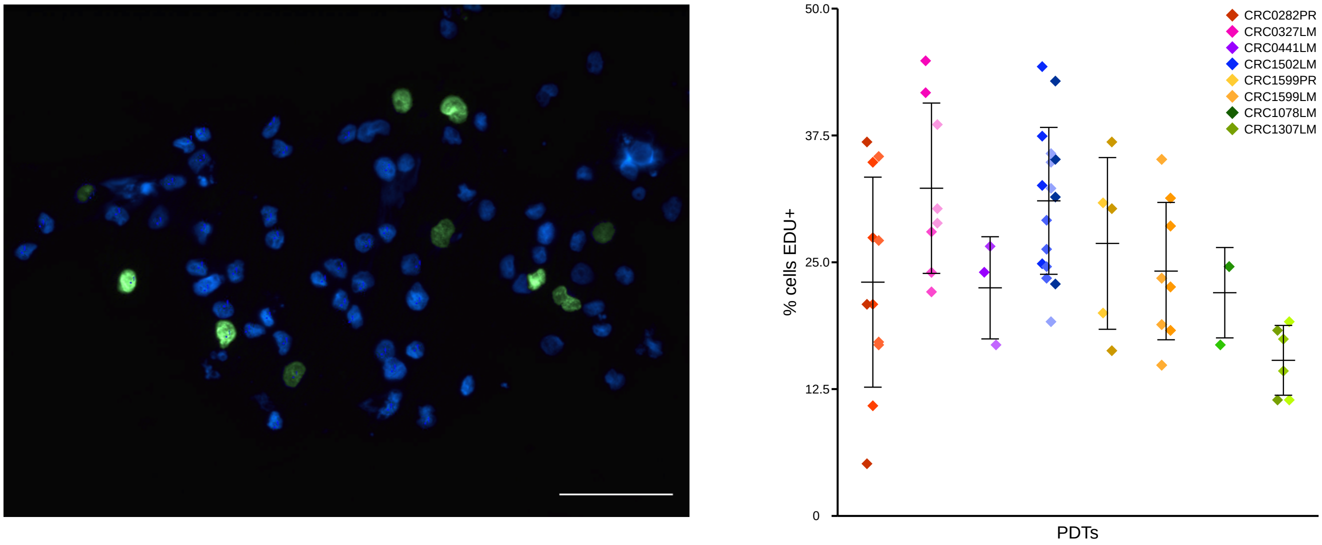


**Fig. S1. EdU staining of PDTs used in MA experiments.**
**a,** Representative EdU staining (green) in the parental CRC1307LM PDT; nuclei (blue) are stained with DAPI (20x magnification, scale bar = 100 μm). **b,** Percentage of EdU positive cells in PDT clones during the MA experiment. Each dot represents the percentage of EdU-positive cells detected in an individual slide (total slides analysed, *N* = 68). Colors correspond to the parental PDT (*N* = 8), while shades of colors indicate the T0 clone from which the cells were derived (*N* = 3-16 for each PDT; total *N* = 22). Error bars indicate the average ±1 standard deviation.

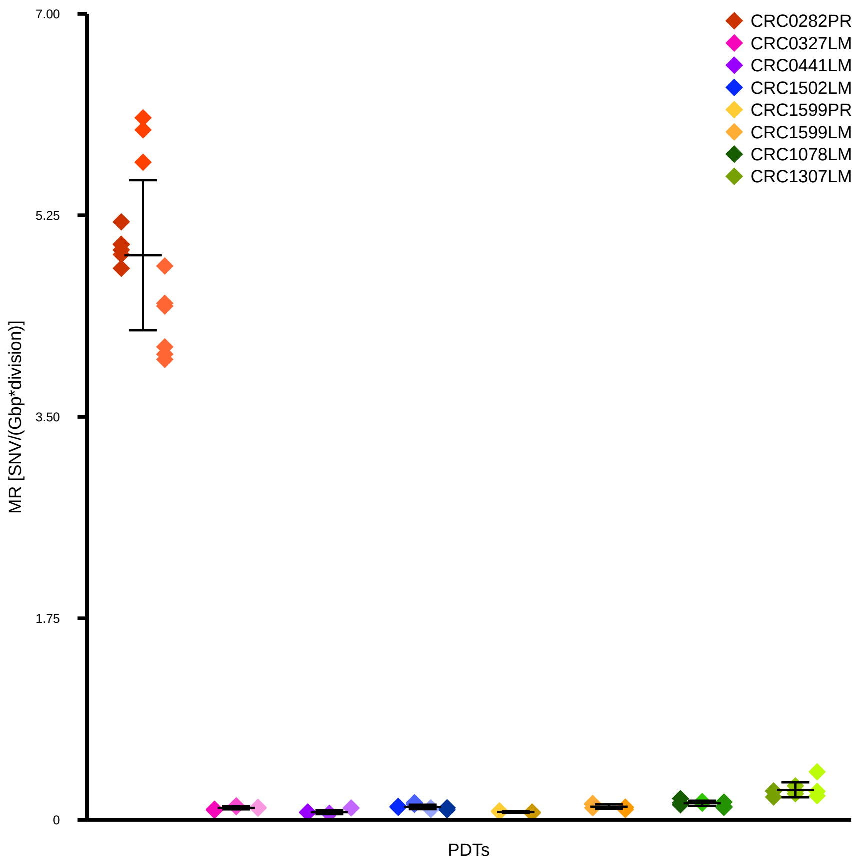


**Fig. S2. Indel MRs in PDTs.**
Each dot represents the MR estimate of individual T1 subclones (*N* = 6-15 for each PDT; total *N* = 73). Colors correspond to the parental PDT (*N* = 8), while color shades indicate the T0 clone from which each T1 subclone originated. Error bars indicate the average ±1 standard deviation.

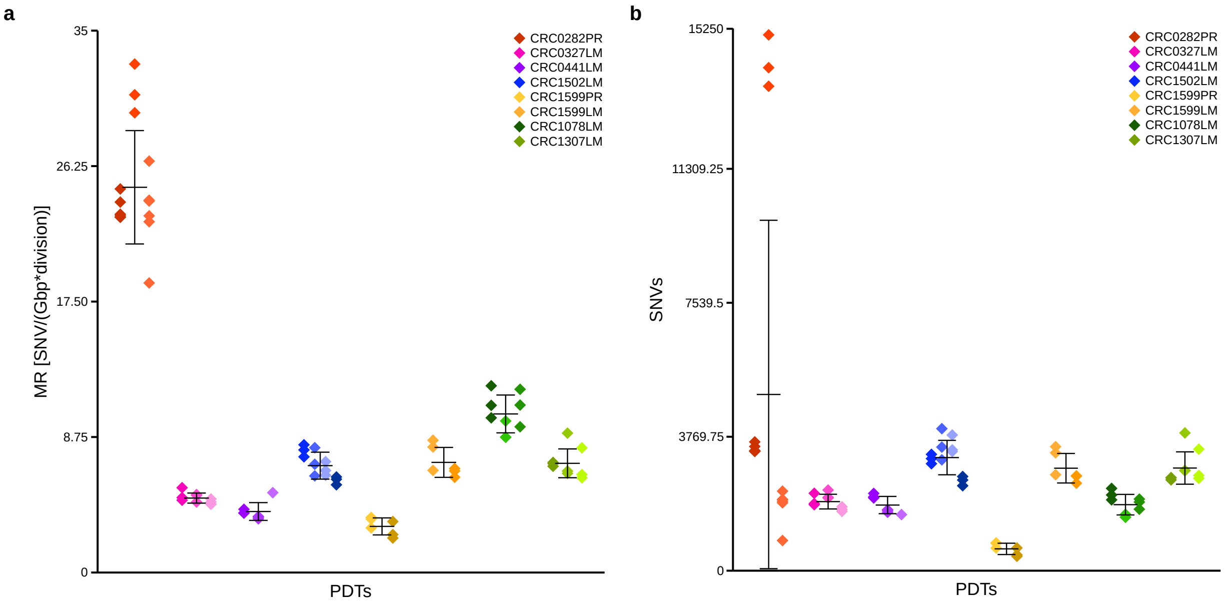


**Fig. S3. MRs calculated based on cell-population doublings and absolute number of accumulated SNVs.**

**a,** SNV MRs based on cell-population doublings in PDTs. Each dot represents the MR estimate of individual T1 subclones (*N* = 6-15 for each PDT; total *N* = 73). Colors correspond to the parental PDT (*N* = 8), while color shades indicate the T0 clone from which each T1 subclone originated. Error bars indicate the average ±1 standard deviation. Spearman correlation between averages shown here and the average MR for each PDT (calculated based on EdU-measured DNA replications) shown in Figure 1, 0.9 (*P* = 0.005). **b,** Absolute number of accumulated SNVs in PDTs. Each dot represents the number of accumulated mutations in individual T1 subclones (*N* = 6-15 for each PDT; total *N* = 73). Colors correspond to the parental PDT (*N* = 8), while color shades indicate the T0 clone from which each T1 subclone originated . Error bars indicate the average ±1 standard deviation. Spearman correlation between averages shown here and the average MR of each PDT shown in Figure 1, 0.71 (*P* = 0.06).

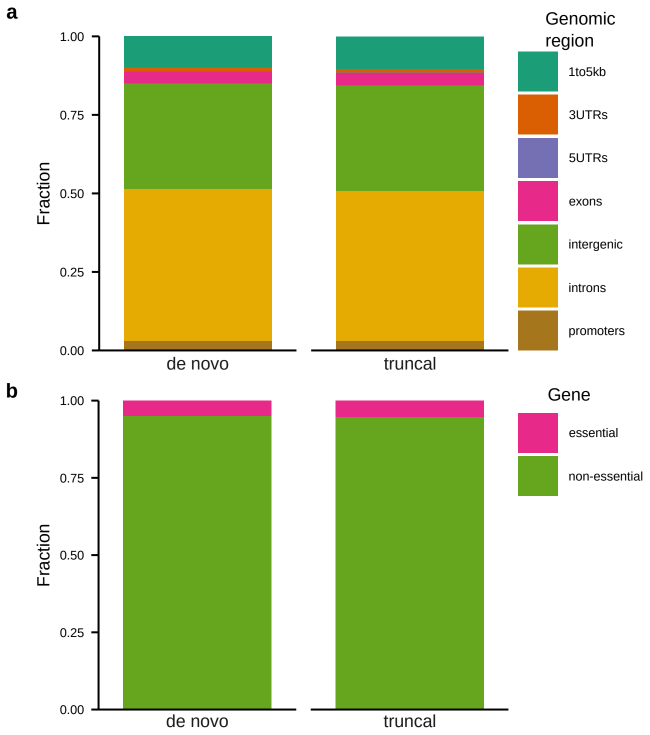


**Fig. S4. Genomic context of *de novo* accumulated SNVs and truncal variants.**
**a,** Genomic localization of the accumulated SNVs detected in all T1 subclones (*de novo*; $N$ = 125,842) compared with that of SNVs detected in parental PDTs (truncal; $N$ = 236,462). Chi-squared test, $P$ = 4.9e-6. **b,** SNVs in essential and non-essential genes accumulated in all T1 clones (*de novo*; $N$ = 753) or detected in the parental PDTs (truncal; $N$ = 1,539). Fisher test, $P$ = 0.84 (odds ratio, 0.94).

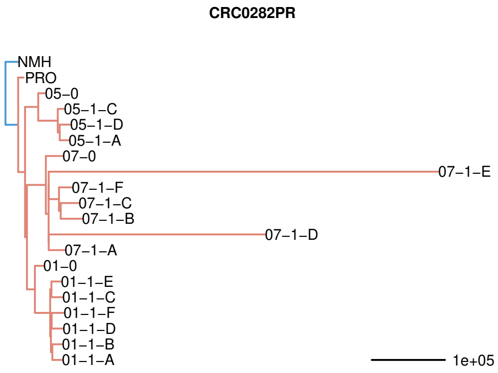
CRC0282PR
NMH
PRO
05–0
05–1–C
05–1–D
05–1–A
07–0
07–1–E
07–1–F
07–1–C
07–1–B
07–1–D
07–1–A
01–0
01–1–E
01–1–C
01–1–F
01–1–D
01–1–B
01–1–A
1e+05

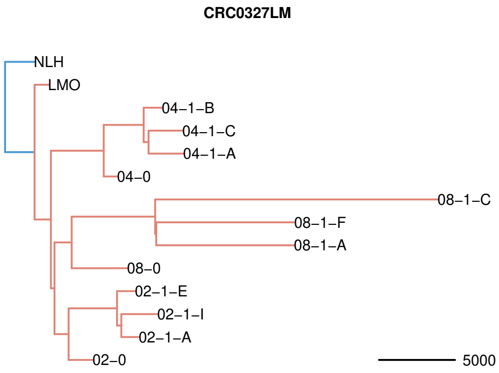
CRC0327LM
NLH
LMO
04–1–B
04–1–C
04–1–A
04–0
08–1–C
08–1–F
08–1–A
08–0
02–1–E
02–1–I
02–1–A
02–0
5000

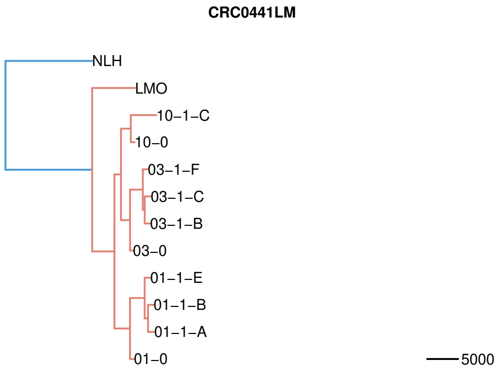
CRC0441LM
NLH
LMO
10–1–C
10–0
03–1–F
03–1–C
03–1–B
03–0
01–1–E
01–1–B
01–1–A
01–0
5000

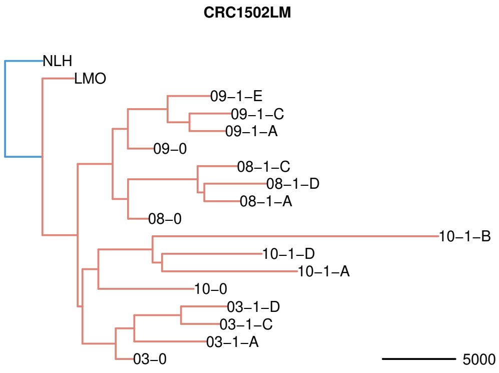
CRC1502LM
NLH
LMO
09–1–E
09–1–C
09–1–A
09–0
08–1–C
08–1–D
08–1–A
08–0
10–1–B
10–1–D
10–1–A
10–0
03–1–D
03–1–C
03–1–A
03–0
5000

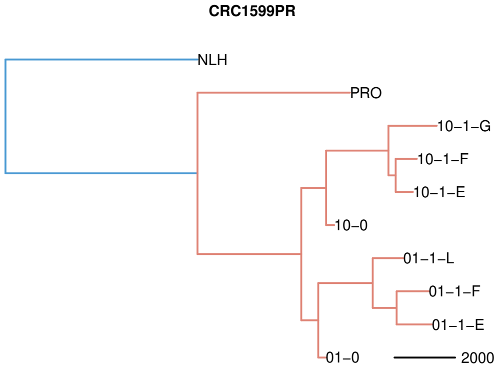
CRC1599PR
NLH
PRO
10–1–G
10–1–F
10–1–E
10–0
01–1–L
01–1–F
01–1–E
01–0
2000

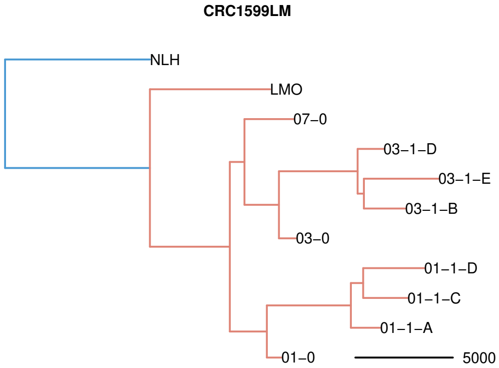
CRC1599LM
NLH
LMO
07–0
03–1–D
03–1–E
03–1–B
03–0
01–1–D
01–1–C
01–1–A
01–0
5000

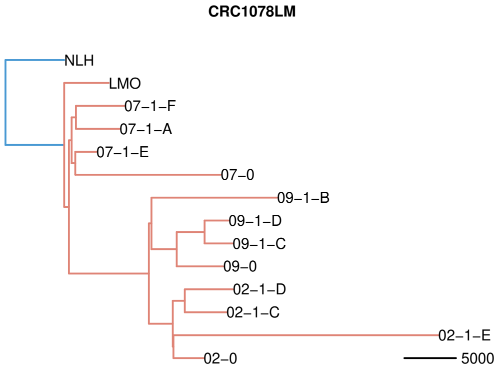
CRC1078LM
NLH
LMO
07–1–F
07–1–A
07–1–E
07–0
09–1–B
09–1–D
09–1–C
09–0
02–1–D
02–1–C
02–1–E
02–0
5000

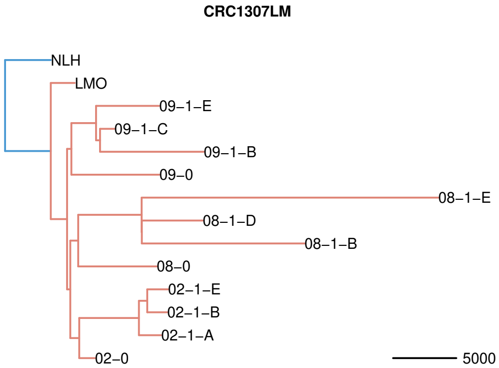
CRC1307LM
NLH
LMO
09–1–E
09–1–C
09–1–B
09–0
08–1–E
08–1–D
08–1–B
08–0
02–1–E
02–1–B
02–1–A
02–0
5000

**Fig. S5. Phylogenetic trees inferred from SNVs.**

Phylogenetic reconstruction of the relationship between clones used in MA experiments, their parental PDTs (LMO or PRO), and the corresponding germline tissues (NLH or NMH). The scale bar indicates the inferred branch lengths. The structure of clone IDs is described in the methods; LMO, PDTs from liver metastases; PRO, PDTs from primary tumors; NLH, normal liver; NMH, normal intestinal mucosa adjacent to the primary tumor.

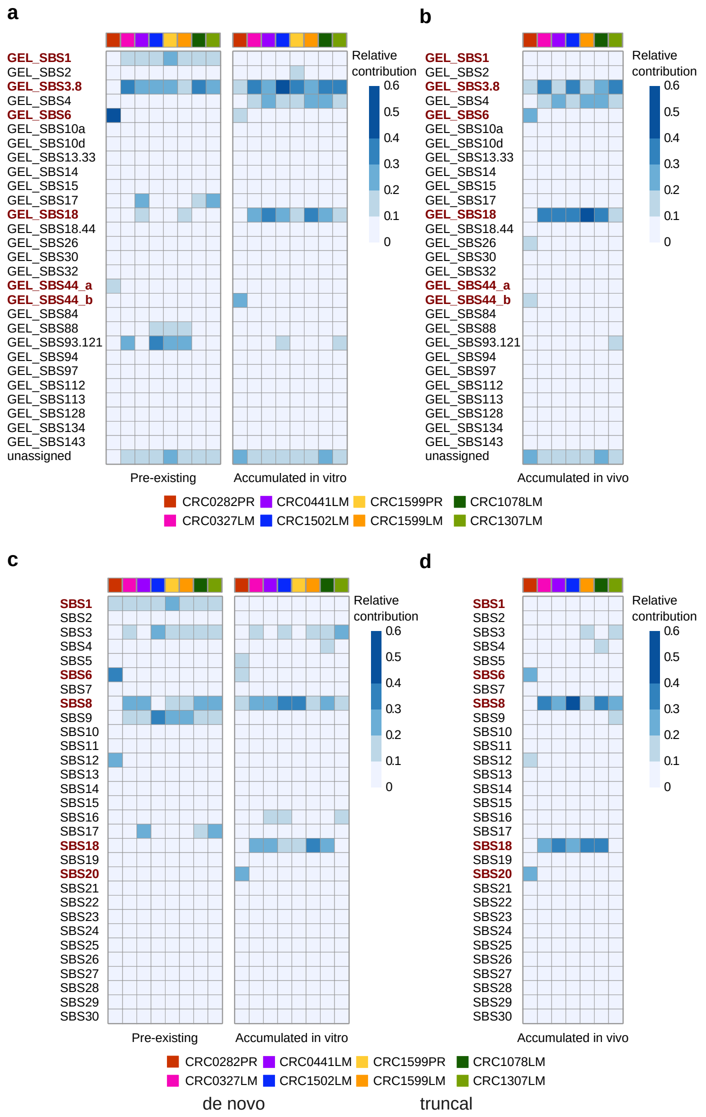


**Fig. S6. Signal- and sigfit-based identification of mutational signatures contributing to MA *in vitro* and *in vivo*.**

**a-b,** Relative contribution of CRC signatures from the GEL dataset (Genomics England), as inferred by the Signal tool[24], to in vitro (**a**) and *in vivo* (**b**) MA. MSI-related signatures (GEL_SBS6 and GEL_SBS44_a and _b) and signatures that changed during the MA experiment (GEL_SBS1, GEL_SBS3.8 and GEL_SBS18) are highlighted in red. **c-d**, Relative contribution of COSMIC v2 signatures, as inferred by the sigfit tool[68], to *in vitro* (**c**) and *in vivo* (**d**) MA. MSI-related signatures (SBS6 and SBS20) and signatures arising during the MA experiment (SBS1, SBS8 and SBS18) are highlighted in red.

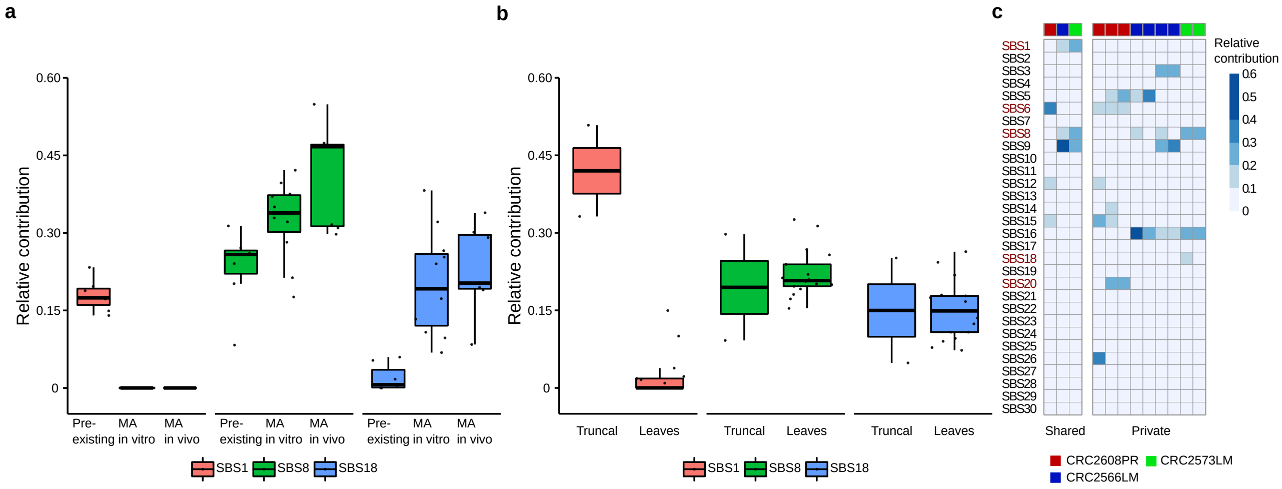


**Fig. S7. Relative contribution of SBS1, SBS8 and SBS18 mutational signatures to pre-existing truncal mutations and to alterations acquired *de novo* during the MA experiment or private to individual clones in patients.**

**a,** Relative contribution of SBS1, SBS8 and SBS18 to the SNVs detected in parental PDTs (Pre-existing), compared to those accumulated during the MA experiment in PDTs (MA in vitro) or in PDXs (MA in vivo) for MSS tumors. Pre-existing vs MA in vitro: $P$ = 9e-5 (SBS1), $P$ = 0.015 (SBS8), $P$ = 5.7e-4 (SBS18), two-tailed Wilcoxon rank sum test. Pre-existing vs MA in vivo: $P$ = 0.001 (SBS1), $P$ = 0.002 (SBS8), and $P$ = 0.002 (SBS18), two-tailed Wilcoxon rank sum test. **b,** Relative contribution of SBS1, SBS8 and SBS18 in truncal SNVs (Truncal) and in SNVs private to individual clones (Leaves) isolated from MSS CRCs in Roerink et al[20]. Truncal vs Leaves: $P$ = 0.02 (SBS1) $P$ = 0.84 (SBS8), and $P$ = 0.95 (SBS18), two-tailed Wilcoxon rank sum test. Each dot indicates the relative contribution of a signature in a specific tumor; boxplots represent the overall distribution of the relative contributions, with default thresholds for whiskers. **c,** Relative contribution of COSMIC signatures to the SNVs shared by different clones derived from the same tumor or private to individual clones in one MSI CRC (CRC2608PR) and two MSS CRCs (CRC2573LM and CRC2566LM). MSI-related signatures (SBS6 and SBS20) and signatures that changed during the MA experiment (SBS1, SBS8 and SBS18) are highlighted in red.

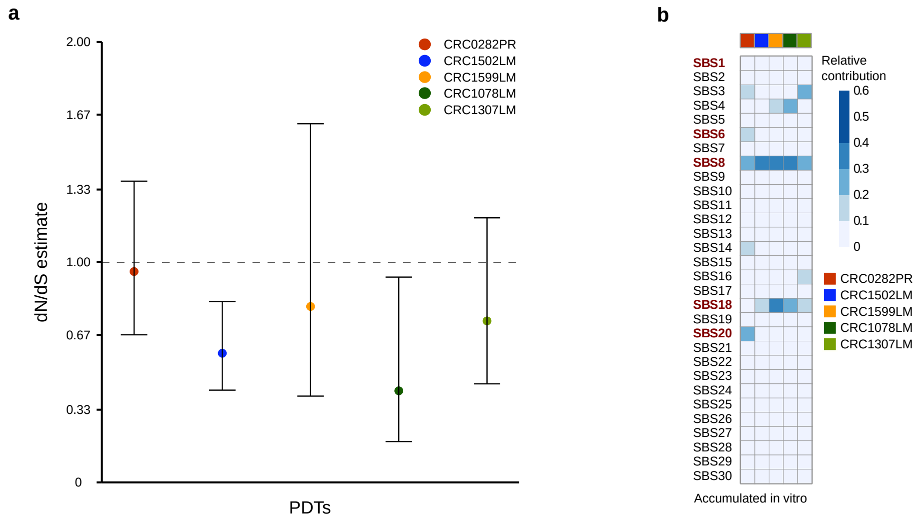


**Fig. S8. Characterisation of *de novo* mutations accumulated during the second round of MA**
**a,** dN/dS estimates for the whole set of *de novo* variants accumulated during the second round of MA experiments (months 7-12). The maximum likelihood estimate of each PDT, obtained using the statistical model dNdScv, is represented as a colored dot. Error bars indicate the upper and lower bounds of the inferred dN/dS values. **b,** Relative contribution of COSMIC v2 signatures (SBS) to the SNVs acquired *de novo* in T2 clones. MSI-related signatures (SBS6 and SBS20) and signatures that changed during the MA experiment (SBS1, SBS8 and SBS18) are highlighted in red.

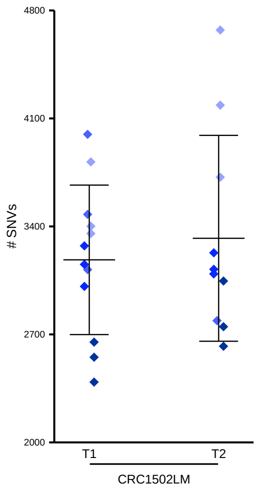


**Fig. S9. Absolute number of accumulated *de novo* SNVs in CRC1502LM clones.**
Each dot represents the total number of accumulated SNVs in T1 subclones ($N$ = 12) or T2 subclones ($N$ = 10); Shades of colors indicate the parental clone from which each T1 or T2 subclone originated. Error bars represent the average ±1 standard deviation.

**CRC0282PR**

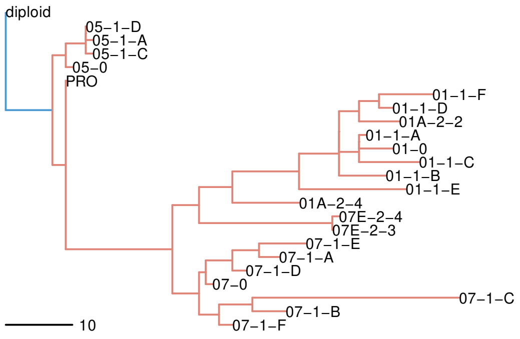


**CRC0327LM**

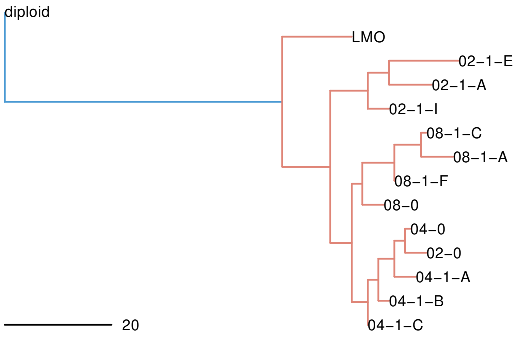


**CRC0441LM**

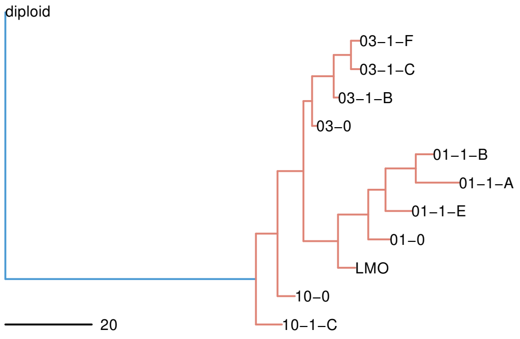


**CRC1502LM**

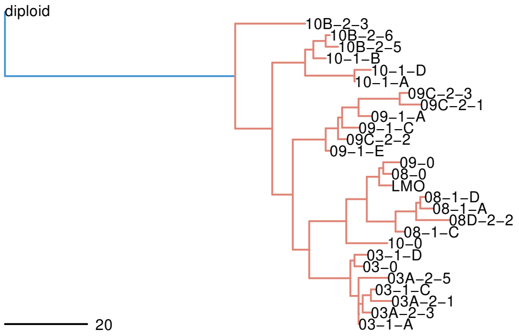


**CRC1599PR**

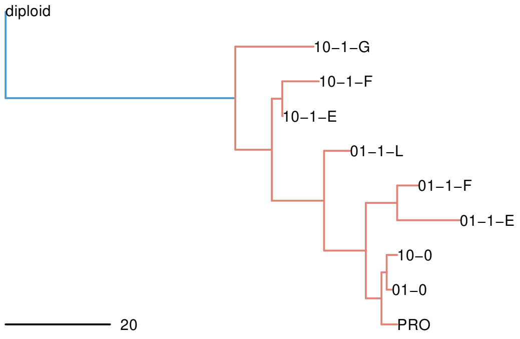


**CRC1599LM**

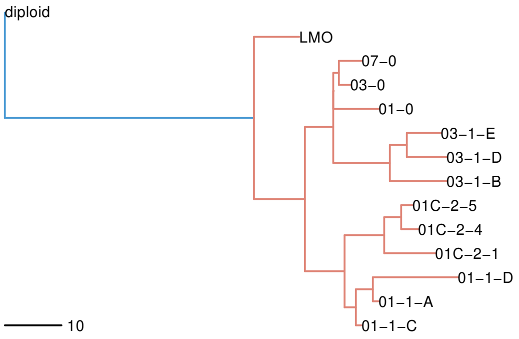


**CRC1078LM**

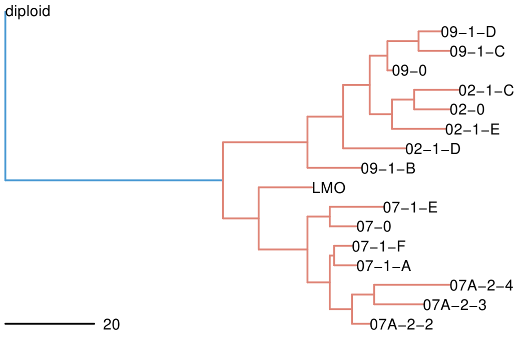


**CRC1307LM**

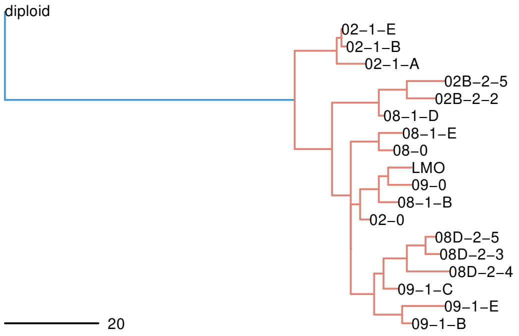

**Fig. S10. Phylogenetic trees inferred from MEDICC2 CN events.**
Phylogenetic reconstruction of the relationship between clones used in MA experiments and their parental PDTs (LMO or PRO) using CNAs derived from haplotype phasing prediction and MEDICC2 (Methods). The scale bar indicates the branch lengths. All trees are rooted on a synthetic diploid genome. The structure of clone IDs is described in the methods; LMO, PDTs from liver metastases; PRO, PDTs from primary tumors; NLH, normal liver; NMH, normal intestinal mucosa adjacent to the primary tumor.

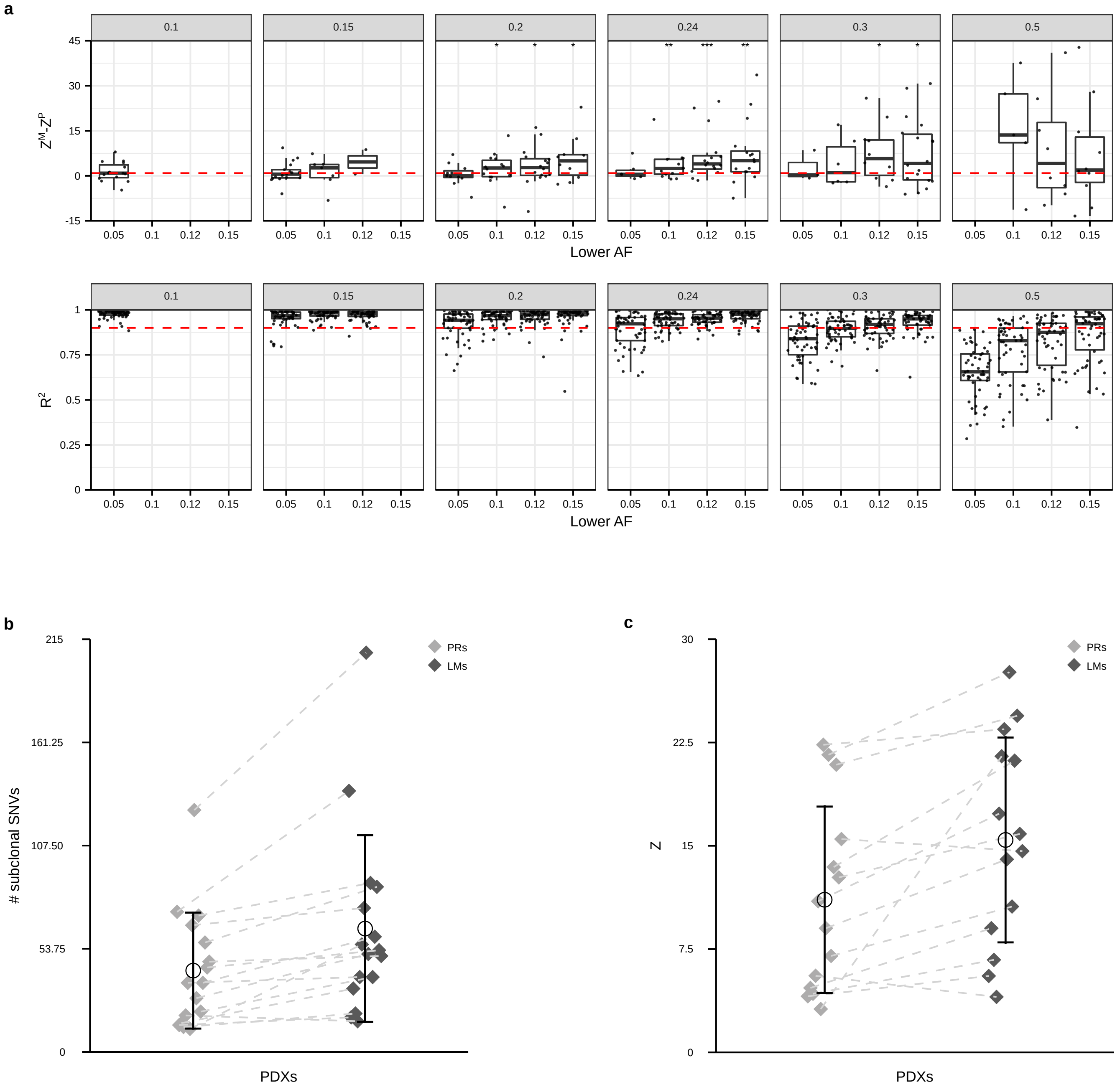
a
0.1
0.15
0.2
0.24
0.3
0.5
$Z^M$-$Z^P$
45
30
15
0
-15
0.05
0.1
0.12
0.15
Lower AF
$R^2$
1
0.75
0.5
0.25
0
b
### subclonal SNVs
215
161.25
107.50
53.75
PRs
LMs
PDXs
c
Z
30
22.5
15
7.5

**Fig. S11. Robustness of MR inferences in the cohort of synchronous primary tumors and metastases.**

**a,** Upper panels: differences ($Z^M$-$Z^P$), for each VAF threshold, between MR estimates of PDXs derived from liver metastases ($Z^M$) and MR estimates of PDXs derived from matched synchronous resections of primary CRCs ($Z^P$). Lower bounds to define subclonal VAFs are indicated on the x axis, while upper bounds are reported as sub-panel titles. For each VAF threshold, only primary-metastasis pairs meeting QC criteria ($R^2$ > 0.9 and > 10 subclonal mutations) are shown, each dot corresponds to a pair; box plots represent the overall distribution of differences. The horizontal dashed red line corresponds to 0; asterisks indicate the results of paired Wilcoxon signed rank tests (* *P* < 0.05, ** *P* < 0.01, *** *P* < 0.001). Bottom panels: $R^2$ values obtained by applying different VAF thresholds to identify subclonal mutations; lower bounds are indicated on the x axis, while upper bounds are reported as sub-panel titles. Each dot corresponds to a tumor; box plots represent the overall distribution of $R^2$ values. The horizontal dashed red line indicates the $R^2$ threshold (0.9) adopted to define good-quality fits. **b,** Total number of subclonal mutations (0.12 < VAF < 0.24) in PDXs derived from matched synchronous resections of primary and metastatic tumors (*N* = 16). Dashed lines connect primary tumors (PRs) to their corresponding liver metastases (LMs). Error bars indicate the average ±1 standard deviation; the average is represented by an empty circle. Paired Wilcoxon signed rank test *P* = 0.0003. **c,** Numerical values of the slopes for the linear regression between cumulative frequency distribution and inverse frequency (Z) of subclonal mutations (0.12 < VAF < 0.24) in paired primary and metastatic CRC PDXs (*N* = 14), after excluding the two PDX pairs showing the largest difference. Dashed lines connect primary tumors (PRs) to their corresponding liver metastases (LMs). *P* = 0.0006, paired Wilcoxon signed rank test. Error bars show the average ±1 standard deviation; the average is represented by an empty circle.

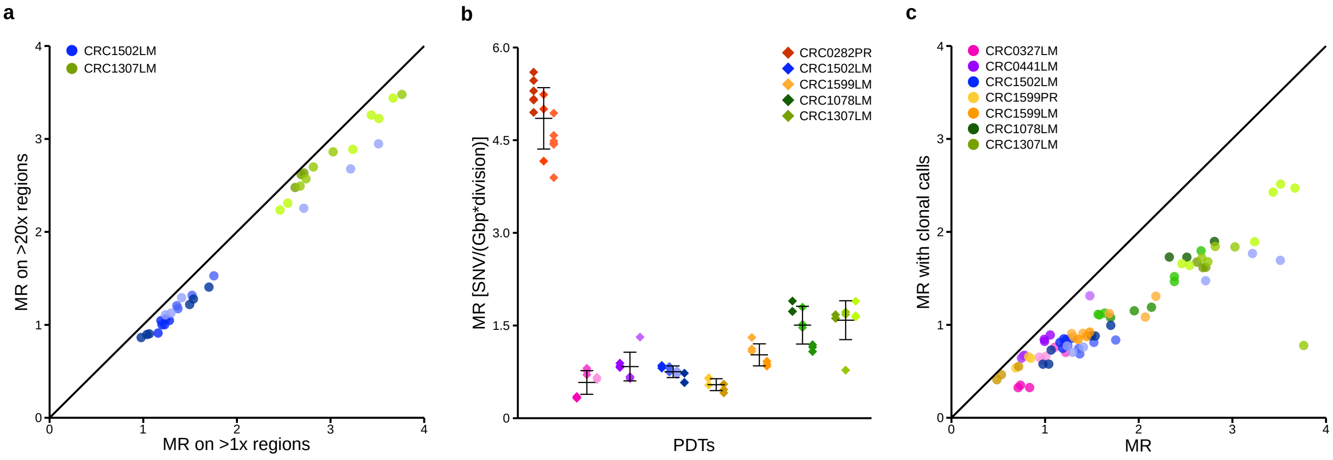


**Fig. S12. Results of alternative approaches for MR computation.**
**a,** Correlation between MR estimates obtained by filtering for genomic regions with coverage > 1x (x axis) and those computed only on regions with a coverage of at least 20x (y axis) for 2 PDTs. Each dot represents a clone (CRC1307LM, $N$ = 14; CRC1502LM, $N$ = 22 ). Spearman correlation 0.98; $P$ < 2e-16 (CRC1307LM) and $P$ < 2.9e-6 ( CRC1502LM). **b,** SNV mutation rates computed by excluding subclonal variants. Each dot represents the MR estimate of individual T1 subclones ($N$ = 6-15 for each PDT; total $N$ = 73); colors correspond to the parental PDT ($N$ = 8), while color shades indicate the T0 clone from which each T1 subclone originated; $P$ = 3e-11, Kruskal-Wallis rank sum test. **c,** Correlation between MR estimates obtained by using the full set of mutations (x axis) and those computed when excluding subclonal variants (y axis). The black line represents the diagonal x=y. Spearman correlation between the two MRs: 0.9; $P$ < 2.2e-16.

**List of Supplementary Tables**

**Table S1. Overview of the main clinical and molecular characteristics of the MA cohort.** Y, yes; N, no; wt, wild type; For each gene column, Mut/CN indicates the type of alteration that has been reported for that specific gene.

**Table S2. List of SNVs detected in parental PDTs and those acquired *de novo* during the MA experiment.** Only SNVs located in regions with CN 1, 2 or 3 are listed; data relative to each of the 8 selected PDTs are reported in individual sheets. For PDTs, VAFs are shown while, for clones, the number of reads supporting each SNV and the total coverage of that genomic position are reported. Coordinate hg38, 1 based hg38 coordinate for the SNV; Ref seq, Reference allele nucleotide; Alt seq, Mutated allele nucleotide; Genomic context, Genomic context for the SNV as reported by Annovar (https://annovar.openbioinformatics.org/en/latest/user-guide/gene/); Gene, Gene that harbors the SNV or nearest one (for intergenic SNVs, the nearest gene to the alteration); Effect, Consequence of the SNVs for exonic ones, . otherwise; Protein change, Effect on the protein for non synonymous SNVs, . otherwise; Essential gene, Essential or non-essential, based on [21].

**Table S3. MRs of individual T1 and T2 clones and their related measurements.** Sequencing_id, original raw data ID the sample, with the nomenclature described in Material and Methods; MR_SNVs, gained SNVs / (CN corrected genome length * total divisions estimated by EdU); MR_indels, gained indels / (CN corrected genome length * total divisions estimated by EdU); N_SNVs, absolute number of gained SNVs; N_indels, absolute number of gained indels; Genome_Length, CN corrected genome length; Generations_EDU, total divisions estimated by EdU staining; Generations_cell_doublings, total divisions estimated by cell population doublings.

**Table S4. Relative contribution of COSMIC v2 signatures (SBS) to the SNVs detected in parental PDTs (Pre-existing), to those acquired *de novo* during the MA experiment in PDTs (Accumulated in vitro) and to the ones acquired *de novo* during *in vivo* MA (Accumulated in vivo).** The 'cosine' sheet reports cosine similarities between the original mutational profile and the one resulting from the fit. vitroMA, in vitro MA; vivoMA, in vivo MA; parental, parental PDTs.

**Table S5. Mutational signatures' contribution to mutation accumulation *in vitro* and *in vivo*, based on alternative computational methods and different signature collections.** The 'Signal' sheet reports contributions of CRC signatures from the GeL dataset (Genomic EngLands), as inferred by the Signal tool[24]. The 'sigfit' sheet reports contributions of COSMIC v2 signatures, as inferred by the sigfit tool[68]. vitroMA, in vitro MA; vivoMA, in vivo MA; parental, parental PDTs.

**Table S6. Clinical and molecular characteristics and mutational profiles of PDTs cloned directly from tumors at the time of surgery**. The 'summary' sheet reports the main clinical and molecular characteristics of this cohort, while sheets named by PDTs' identity contain the list of SNVs detected in each model. Only SNVs located in regions with CN 1, 2 or 3 are listed. For individual clones, the number of reads supporting each SNV and the total coverage of that genomic position is reported, while for the shared SNVs the average number of supporting reads and total coverage are shown. Y, yes; N, no; wt, wild type; For each gene column, Mut/CN indicates the type of alteration that has been reported for that specific gene. Coordinate hg38, 1 based hg38 coordinate for the SNV; Ref seq, Reference allele nucleotide; Alt seq, Mutated allele nucleotide; Genomic context, Genomic context for the SNV reported by Annovar (https://annovar.openbioinformatics.org/en/latest/user-guide/gene/); Gene, Gene that harbors the SNV or nearest one (for intergenic SNVs, the nearest gene to the alteration); Effect, Conseguence of the SNVs for exonic ones, . otherwise; Protein change, Effect on the protein for non synonymous SNVs, . otherwise; Essential gene, Essential or non-essential, based on [21].

**Table S7. Relative contribution of COSMIC v2 signatures to private and shared SNVs in clonal PDTs directly derived from patient material.** The 'cosine' sheet reports cosine similarities between the original mutational profile and the one resulting from the fit. Private, derived from SNVs detected in individual clones; shared, derived from SNVs shared by multiple clones derived from the same patient; leaves, derived from SNVs assigned to the leaves in phylogenetic trees from Roerink et al.[20]; truncal, derived from SNVs assigned to the tumor MRCA node in phylogenetic trees from Roerink et al[20].

**Table S8. Relative contribution of COSMIC v2 signatures to the SNVs acquired *de novo* during MA in months 7-12.** The 'cosine' sheet reports cosine similarities between the complete profile and the one resulting from the fit. Only SNVs found in regions with CN 1, 2 or 3 are listed. The set of data relative to each of the 5 original tumors for which T2 clones were available is reported in an individual sheet; for each mutation the number of reads supporting the SNVs and the total coverage of that specific genomic location are shown. Coordinate hg38, 1 based hg38 coordinate for the SNV; Ref seq, Reference allele nucleotide; Alt seq, Mutated allele nucleotide; Genomic context, Genomic context for the SNV reported by Annovar (https://annovar.openbioinformatics.org/en/latest/user-guide/gene/); Gene, Gene that harbors the SNV or nearest one (for intergenic SNVs); Effect, Conseguence of the SNVs for exonic ones, . otherwise; Protein change, Effect on the protein for non synonymous SNVs, . otherwise; Essential gene, Essential or non-essential, based on [21].

**Table S9. Non synonymous alterations common to all CRC1502-T2 hypermutating clones and analysis of the total mutational burden in cases with or without mutations in those genes, in TCGA and in a proprietary PDX collection**. The 'Shared alterations' sheet lists the non synonymous alterations, the 'WES' and 'TCGA' sheets shows the results of one-tailed Wilcoxon rank sum test comparing cases with or without mutations in the two dataset, respectively. Coordinate hg38, hg38 1 based coordinate; Ref seq, Reference allele nucleotide; Alt Seq, Mutated allele nucleotide; status, inherited for SNVs already present in the T1 ancestor, denovo_private for those only shared by all the T2 hypermutating clones; median_mut, median total tumor burden in mutated samples; median_wt, median total tumor burden in WT samples; n_mut, n of mutated samples; n_w, n of WT samples; W, Wilcoxon rank sum test statistic; padj, Benjamini and Hochberg multiple test corrected p-value

**Table S10. Subclonal SNV counts in clones originated from CRC1078LM, CRC1307LM, CRC1599PR and CRC1599LM.** Sequencing_id, original raw data ID; n_subclonal, number of subclonal muts.

**Table S11. Mutational counts and fitting of the subclonal variants' distribution based on different subclonal-VAF thresholds in WES data from PDXs that originated from matched Primary-Metastatic tumor pairs.** Sample_id, human readable ID; Sequencing_id, original raw data ID; lower_VAF, Lower threshold used to identify subclonal mutations; upper_VAF, Higher threshold used to identify subclonal mutations; R2, R2 of the fit on the cumulative number of subclonal mutations; slope, slope of the previous fit; n_subclonal, Number of subclonal mutations identified; n_total, Total number of mutations.

**Table S12. Relevant metrics obtained from picard wgsmetrics relative to all the 30x WGSs generated from MA clones, bulk organoids and their matched normal tissue.** Sample_id, human readable ID; Sequencing_id, original raw data ID; mean_coverage, The mean coverage in bases of the genome territory, after all filters are applied; sd_coverage, The standard deviation of coverage of the genome after all filters are applied; median_coverage, The median coverage in bases of the genome territory, after all filters are applied.

**Table S13. Results of GATK CalculateContamination relative to all the WGSs 30x for which mutation calling was performed.** Sequencing_id, original raw data ID; contamination, fraction of reads due to cross-sample contamination for each tumor sample.

**Table S14. Relevant metrics obtained from picard wgsmetrics and GATK Contamination estimates relative to WGSs from clonal PDTs that were directly derived from patient material.** The 'WGS QC' sheet reports picard wgsmetrics results, the 'contamination' sheet the contamination estimates. Sample_id, human readable ID; Sequencing_id, original raw data ID; mean_coverage, The mean coverage in bases of the genome territory, after all filters are applied; sd_coverage, The standard deviation of coverage of the genome after all filters are applied; median_coverage, The median coverage in bases of the genome territory, after all filters are applied; contamination, fraction of reads due to cross-sample contamination for each tumor sample.

**Table S15. Relevant metrics obtained from picard hsmetrics relative to WES data from PDXs that originated from matched Primary-Metastatic tumor pairs.** Sequencing_id, original raw data ID; mean_target_coverage, The mean coverage of a target region; median_target_coverage, The median coverage of a target region.
